# pan-MHC and cross-Species Prediction of T Cell Receptor-Antigen Binding

**DOI:** 10.1101/2023.12.01.569599

**Authors:** Yi Han, Yuqiu Yang, Yanhua Tian, Farjana J. Fattah, Mitchell S. von Itzstein, Yifei Hu, Minying Zhang, Xiongbin Kang, Donghan M. Yang, Jialiang Liu, Yaming Xue, Chaoying Liang, Indu Raman, Chengsong Zhu, Olivia Xiao, Jonathan E. Dowell, Jade Homsi, Sawsan Rashdan, Shengjie Yang, Mary E. Gwin, David Hsiehchen, Yvonne Gloria-McCutchen, Ke Pan, Fangjiang Wu, Don Gibbons, Xinlei Wang, Cassian Yee, Junzhou Huang, Alexandre Reuben, Chao Cheng, Jianjun Zhang, David E. Gerber, Tao Wang

## Abstract

Profiling the binding of T cell receptors (TCRs) of T cells to antigenic peptides presented by MHC proteins is one of the most important unsolved problems in modern immunology. Experimental methods to probe TCR-antigen interactions are slow, labor-intensive, costly, and yield moderate throughput. To address this problem, we developed pMTnet-omni, an Artificial Intelligence (AI) system based on hybrid protein sequence and structure information, to predict the pairing of TCRs of αβ T cells with peptide-MHC complexes (pMHCs). pMTnet-omni is capable of handling peptides presented by both class I and II pMHCs, and capable of handling both human and mouse TCR-pMHC pairs, through information sharing enabled this hybrid design. pMTnet-omni achieves a high overall Area Under the Curve of Receiver Operator Characteristics (AUROC) of 0.888, which surpasses competing tools by a large margin. We showed that pMTnet-omni can distinguish binding affinity of TCRs with similar sequences. Across a range of datasets from various biological contexts, pMTnet-omni characterized the longitudinal evolution and spatial heterogeneity of TCR-pMHC interactions and their functional impact. We successfully developed a biomarker based on pMTnet-omni for predicting immune-related adverse events of immune checkpoint inhibitor (ICI) treatment in a cohort of 57 ICI-treated patients. pMTnet-omni represents a major advance towards developing a clinically usable AI system for TCR-pMHC pairing prediction that can aid the design and implementation of TCR-based immunotherapeutics.

## INTRODUCTION

TCRs bind to antigenic peptides presented by major histocompatibility complexes (MHCs), forming a complex called pMHC on the surface of antigen-presenting cells, which serve as recognition markers for cytotoxic T cells. The fast, cost-effective, and accurate identification of the binding between TCRs and T cell antigens will vastly enhance our understanding of T cells’ roles in normal developmental processes and multiple diseases. Such information will also widely impact the design and implementation of various immunotherapies and disease diagnostics^1–5^, such as engineering of TCR T therapies and TCR-like drugs, antigen selection for tumor vaccines, and biomarkers for monitoring T cell function.

A number of experimental approaches were developed to quantify T cell-antigen recognition, including tetramer assays, TetTCR-seq^6,7^, T-Scan^8^, BEAM-T (10X Genomics), *etc*. However, these approaches remain time-consuming, technically challenging, costly, and still yield only moderate throughput. These deficiencies call for the development of state-of-the-art *in silico* algorithms to predict TCR binding specificity towards T cell antigens. Several studies have generated evidence showing that it is possible to learn the binding between TCRs and T cell antigens *via* machine learning approaches. These include pMTnet, generated previously by us^9,10^, and other tools, such as TCRGP^11^, TCRex^12^, NetTCR^13^, *etc*.

Despite these recent efforts, this scientific problem remains far from solved. First, AUCs of most of these works still hover around 0.8 or lower (though this also depends on the specific validation cohorts). Second, most tools are capable of generating predictions for only TCRs of CD8^+^ T cells and/or only human TCR-pMHC pairs, ignoring TCRs of CD4^+^ T cells and TCR-pMHC pairs from other species, which are equally important. Thirdly, many of these tools only consider the β chains or even only the CDR3 regions of the β chains, and/or only the peptide but not the MHC alleles of pMHCs, an incomprehensive approach that may contribute to poor performance to date. Lastly, insufficient efforts have been put into exploring how such predictive models can result in translational value, especially in prospectively generated data cohorts. Clearly, substantial improvements are needed before such AI systems could be useful for clinical application.

Another central limitation of current approaches to the prediction of TCR-pMHC binding is the exclusive focus on protein and peptide sequences^11–17^, without consideration of their structural features. There is only one work to our knowledge^17^, which predicted binding based on structural features but achieved modest performance in their own validation. One obvious drawback of this work is that it only considered TCR-pMHC complexes with known structures, thus severely limiting the size of available training data. Luckily, with the emergence of large deep learning models, such as Alphafold, RoseTTAfold, and ESMFold^18–20^, it is now possible to model and predict new protein structures given only their amino acid sequences. We posit that the consideration of the structural aspect of TCR-pMHC binding, through a design that properly leverages such models, could improve TCR-pMHC pairing predictions. In particular, in the local environment where TCRs and pMHCs interact, TCRs and peptides are relatively flexible, while the MHCs have more rigid structures. Therefore, we decided to adopt a hybrid model that digests MHC information by a structure-based sub-model and digests TCR and peptide information by sequence-based submodels to predict their pairing.

Here we present a hybrid sequence-structure model to predict pairings between TCRs of αβ T cells and pMHCs, for both class I and II pMHCs and for both human and mouse TCR-pMHC pairs. The pan-MHC and cross-species prediction capability is achieved by large-scale information sharing enabled by this hybrid design. Given the omnibus nature of this pMHC-TCR binding prediction neural network, we named our model, pMTnet-omni. Through a series of validations and benchmarking, we show that pMTnet-omni achieves state-of-the-art performance in distinguishing binding *vs.* non-binding and stronger *vs.* weaker binding of TCR-pMHC pairs. pMTnet-omni interpreted the longitudinal evolution and the spatial heterogeneity of TCR repertoire, from the perspective of antigen binding affinity landscape. Our analyses demonstrate the potential of pMTnet-omni for clinically useful applications, such as prediction of irAE of ICI treatment.

## RESULTS

### A Hybrid Sequence-Structure Model for TCR-pMHC Pairing Prediction

pMTnet-omni improves the accuracy of pMHC-TCR binding prediction by incorporating a novel hybrid sequence-structure approach based on our knowledge of the physical and geometric constraints of TCRs, peptides, and MHCs in the local region where they interact. We employed an overall conquer-by-division design. Specifically, we developed a suite of new sequence encoders for each chain of the TCRs, which consist of a V gene encoder and a CDR3 encoder. We encoded pMHCs so that the peptide parts of pMHCs complexes are encoded by a sequence-based encoder and the MHCs are embedded by the structurally-aware ESM2 encoders in ESMFold^20^. These different protein/peptide encoders were separately pre-trained to leverage the vast amounts of “unlabeled” data that have been accumulated by the field. Namely, these “unlabeled” data include TCR sequencing data without known pMHC binding affinity and pMHC data without TCR specificity. Then we assembled these separate encoders into the final TCR-pMHC binding prediction model, which is trained *via* contrastive learning (**Fig. 1a**). To be compatible with this contrastive paradigm during training, pMTnet-omni generates binding predictions for a query TCR-pMHC pair by comparing the query TCR against a large set of species-matched random TCRs, which forms a null hypothesis. The final output is in the form of a rank-percentile score (rank%), where a smaller rank% means a stronger binding of the query TCRs, as compared with the background TCRs, to a certain pMHC. A description of the most critical components of pMTnet-omni is provided below, with more details in the **method section** and **Sup. Files 1** & **2**.

**Fig. 1.**
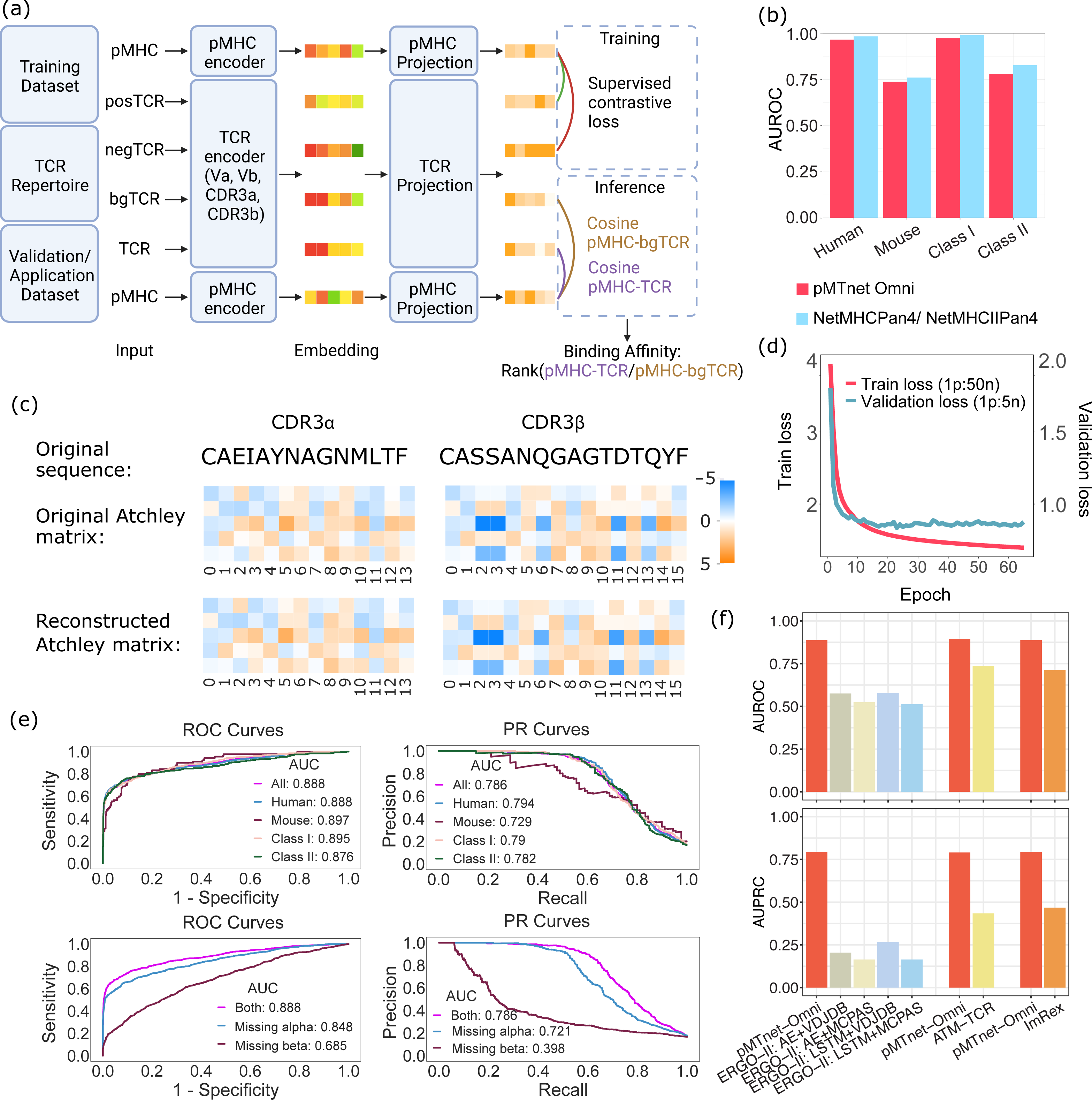
The pMTnet-omni model accurately predicts TCR-pMHC binding. (a) Cartoon diagram of the pMTnet-omni model. posTCR, negTCR, and bgTCR: positive TCR, negative TCR, and background TCR. (b) Validating the p-MHC encoder *via* testing the encoder’s capability to predict peptide *vs* MHC binding. (c) The performance of the CDR3 VAE encoders. The original amino acid sequences, the original Atchley Factor matrices, and the pMTnet-omni’s reconstructed Atchley Factor matrices are shown. (d) The decrease in the loss function over time, for pMTnet-omni. (e) ROC and PR curves showing the performance of pMTnet-omni in the validation dataset. The pMHC-TCR pairs in the validation dataset were classified into four groups according to the species and MHC class: human pairs, mouse pairs, pairs with class I MHC, and pairs with class II MHC. ROC and PR curves of pMTnet-omni were also evaluated on incomplete validation data pairs. (f) AUROCs of pMTnet-omni and six other benchmark software.

We built a peptide-MHC (p-MHC) model to connect the peptide and MHC, and trained this model according to whether or not the peptide and the MHC are binding (**Sup. Fig. 1, Sup. Fig. 2a**). We took this approach because MHCs need to first present peptides on the surface of the antigen-presenting cells before they can be recognized by TCRs. In the p-MHC model, each amino acid of the peptide is converted into five numbers by the Atchely factors^21^, forming an Atchley factor matrix. Then this matrix is digested by a series of attention-based and non-attention-based components. The 1,280-dimensional embeddings generated by ESM-2 for MHC embeddings are fed into our p-MHC model, combined with the peptide embeddings, and processed by a series of layers that output the final peptide-MHC binding prediction. In **Fig. 1b**, we show that this intermediate model achieved very comparable performance to netMHCpan and netMHCIIpan^22,23^, in the validation set. The key innovation in this section of the network is that we enforced large-scale information sharing across class I and II MHCs and across human and mouse pMHC data, by training on all peptide-MHC binding data pooled together and by leveraging the ESM2 encoder. While class I and II pMHCs interact with TCRs in different manners, and human and mouse pMHCs also differ significantly, many features of TCR-pMHC binding are shared across class I and II pMHCs and conserved through vertebrate evolution^24,25^. We assert that pooling all data (I and II, human and mouse) together and training a unified peptide-MHC binding model will enhance the prediction performance of pMTnet-omni overall, and will be particularly helpful for class II and mouse TCR-pMHC pairs with much fewer data points. After this intermediate model was trained, we extracted the last layer before the peptide-MHC binding prediction head, with 30 neurons, to serve as the final peptide-MHC embedding. We then trained TCR encoders (**Sup. Fig. 1, Sup. Fig. 2b**, training data shown in **Sup. Table 1**). An innovative aspect of this step is the creation of four encoders for the Vα, CDR3α, Vβ and CDR3β of TCRs. CDR3s are far more diverse, generated through the “somatic” VDJ recombination process, mainly bind peptides^26^, but feature shorter sequences. In contrast, V gene segments are much less diverse and exclusively germline (without somatic hypermutation), and primarily bind MHCs, but are much longer. These biological characteristics led to our decision to create these four separate encoders. Each of the four encoders was created by a variational auto-encoder (VAE) with attention-based modules, to encode the Atchley factor matrices of the V and CDR3 sequences through a bottleneck layer. As shown in **Fig. 1c** (CDR3s) and **Sup. Fig. 3** (V genes), the reconstructed Atchley factor matrices of CDR3s and V genes are almost identical to the original CDR3s and V genes, evidence of the validity of these TCR encoders. For the V genes, we also visualized the distribution of all possible V genes in their embedding space (**Sup. Fig. 4**). Reassuringly, the different alleles of the same V genes, which usually differ by a limited number of amino acids, are tightly clustered together, again confirming the validity of our encoders.

Finally, we trained a head module to connect all these separate networks to predict pairings between TCRs and pMHCs (**Fig. 1a., Fig. 1d**, training data shown in **Sup. Table 2**). For each binding TCR-pMHC pair in our training dataset, 50 TCRs were randomly sampled from our pooled TCR repertoire (**Sup. Table 1**) to serve as the putative non-binding TCRs for the same pMHC. In pMTnet-omni, the embeddings of these TCRs and the pMHC were projected and normalized to unit hypersphere, where the distance between TCR and pMHC could be measured by dot product. A smaller distance between the truly binding TCR and the pMHC, as compared to the distances between the randomly mismatched TCRs and the pMHC, was pursued during the training process. Consistent with the contrastive learning paradigm during training, the final output of pMTnet-omni was created as a percentile ranking (rank%) score between 0 and 1. Rank% for each query TCR-pMHC pair was computed by comparing the binding of the query TCR-pMHC against a set of 1,000,000 randomly sampled TCRs from the same species (human or mouse) as the query TCR. Rank% closer to 0 represents stronger predicted binding.

To train and validate pMTnet-omni, we curated 168,484 human/mouse class I/II TCR-pMHC pairs from a total of 69 public datasets (**Sup. Table 2**). We employed stringent data curation procedures to curate these data. Details of the datasets as well as the data curation process can be found in **Sup. File 2**. Among the 69 datasets, 19 were used as the training cohort, and the other 50 datasets were used as the independent validation cohort. The training and validation cohorts come from different datasets, and we removed overlapping TCR-pMHC pairs between the training and validation cohorts. During training, the training cohort was further split into a training and an internal validation cohort, for cross-validation purposes. To increase sample size, we took the innovative step of including pairing data with missing data elements in the training cohort, as incomplete records still contain useful information on TCR-pMHC pairing. The numeric embeddings of incomplete data elements were replaced by 0s, during the training process. This approach dramatically increased our training data sample size by three-fold. For the validation cohort, no missingness was allowed, and we ensured none of the pairs in the validation cohort was identical to any pair in the training cohort even if only comparing non-missing data elements of the training pairs.

### Accurate TCR-pMHC Pairing Prediction with pMTnet-omni

In the independent validation cohort, we adopted two metrics to measure the performance of pMTnet-omni: Area Under the Receiver Operating Characteristic Curve (AUROC) and Area Under The Precision-Recall Curve (AUPRC). As is shown in **Fig. 1e**, pMTnet-omni achieved an AUROC of 0.888 and an AUPRC of 0.786 on all validation data including human/mouse and class I/II TCR-pMHC pairs. To evaluate if our knowledge-sharing approach was successful, we separately examined the performance of pMTnet-omni on human vs. mouse TCR-pMHC pairs. As **Fig. 1e** shows, pMTnet-omni performed comparably for human (AUROC=0.888, AUPRC=0.794) and mouse (AUROC=0.897, AUPRC=0.729) pairs. We also evaluated the performance of pMTnet-omni for class I and II pairs. As is shown in **Fig. 1e**, pMTnet-omni also achieved comparable performances for class I (AUROC=0.895, AUPRC=0.79) and class II (AUROC=0.876, AUPRC=0.782) pairs. We also evaluated the relative contributions of the TCR α and the β chains to binding between TCR and pMHCs. For each TCR-pMHC pair in the independent validation cohort, we masked out either the α or β chains and filled embedding with 0s. We then evaluated the performance of pMTnet-omni on these masked datasets. As **Fig. 1e** shows, masking out α chains reduced the AUROC to 0.848 and AUPRC to 0.721, while masking out β chains reduced the AUROC to 0.685 and AUPRC to 0.398, consistent with the role of TCR β chains as the main determinants of TCR-pMHC binding^14^.

We benchmarked pMTnet-omni against other similar software applications, including ERGO-II^26^, ATM-TCR^27^, and ImRex^28^. Each of these software tools has some limitations on what types of TCRs or peptides or MHCs can be used as input. To ensure a fair comparison, in the validation cohort, we only retained the types of TCR-pMHC pairs that are compatible with each of the benchmarked software tools. In this analysis, pMTnet-omni had superior AUROC and AUPRC compared to all other tools (**Fig. 1f**).

Imitating netMHCpan and netMHCIIpan, we suggest two default cutoffs on the rank% scores to call binding TCRs. Strong binders were defined as those with rank%<0.05% (specificity=0.9998, sensitivity=0.2032, F1=0.3374) and weak binders were defined as those with rank%<3% (specificity=0.9933, sensitivity=0.5431, F1=0.689). Unless otherwise stated, we used the weak binding cutoff in this work. We also calculated these performance characteristics for the benchmark software at the same 3% and 0.05% cutoffs on their prediction scores for defining binding *vs* non-binding TCRs (**Sup. Table 3**). pMTnet-omni demonstrated the best overall accuracy.

### Emergence of Tumor Associated Antigen-specific TCR Clones during Tumor Progression

We determined whether pMTnet-omni could depict the evolution of tumor antigen-specific TCR repertoire during tumor progression. The lung adenocarcinoma cell line 344SQ^29^, derived from the metastases of a *Kras*^G12D^/*Tp53*^*R172H*^ genetically engineered mouse model, was subcutaneously injected into seven 129/Sv mice (**Fig. 2a**). The tumor cells metastasized to the lung around the 5^th^/6^th^ week. The primary tumors at the subcutaneous site and the lungs of the mice were harvested from week 3 to week 9 (a different mouse each week). Multiplexed scRNA-seq/scTCR-seq data were generated from the primary tumors and lungs of these 7 mice. We performed cell typing according to gene expression patterns and tumor somatic mutations that can be called from the scRNA-seq reads^30^ (**Fig. 2b**, and more details provided in **Sup. File 3**).

**Fig. 2.**
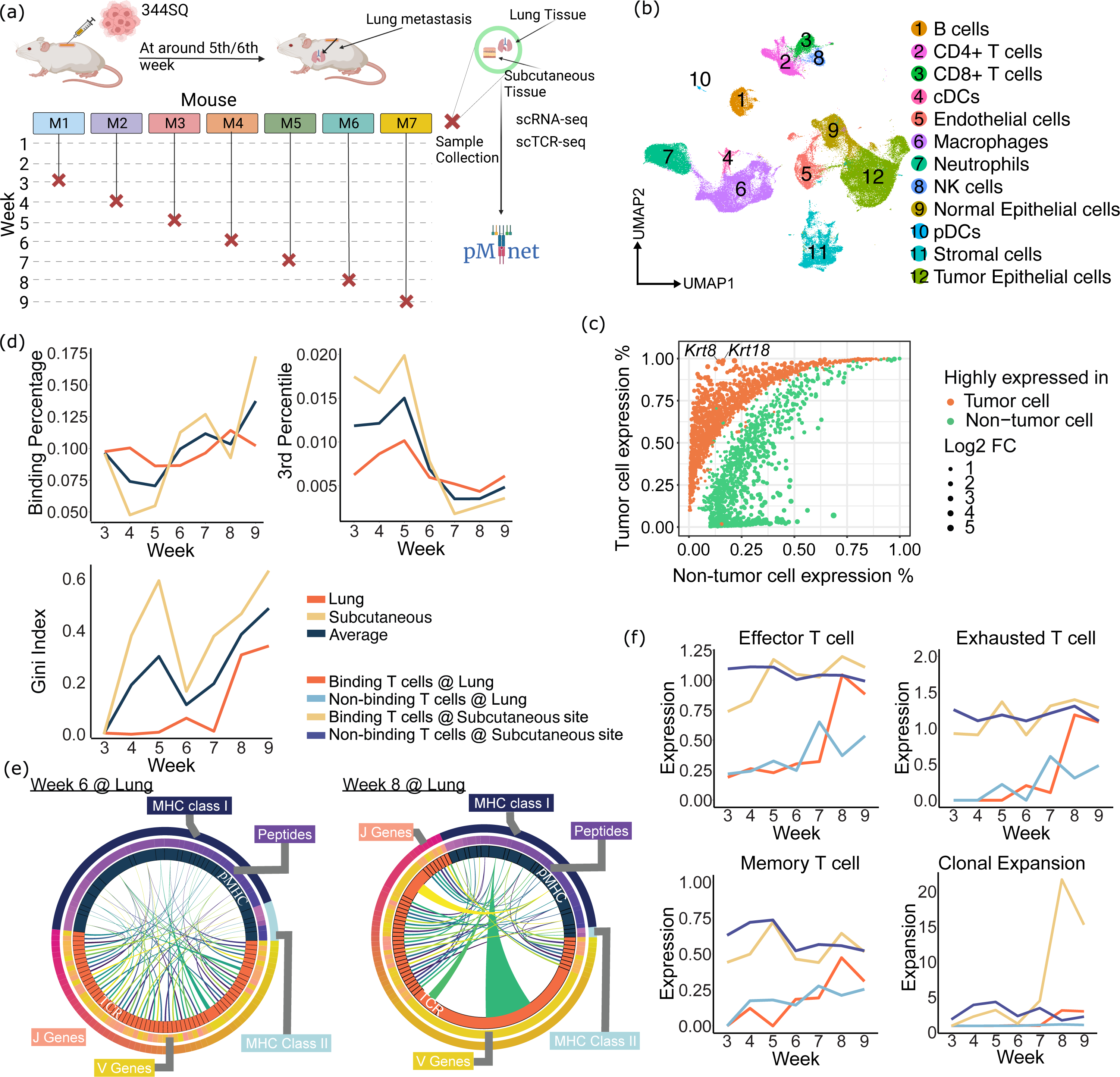
pMTnet-omni interprets TCR evolution in the 344SQ/129Sv mouse model. (a) Schema of the experiment design of mouse model and data collection. (b) UMAP showing cell clustering, of all cells from both the subcutaneous (primary) and lung (metastatic) tissue samples and also from all 7 time points. (c) Differential expression of tumor antigen candidates in the tumor epithelial cells and cells of all other cell types combined. (d) Percentages of predicted binding TCR-pMHC pairs, 3rd percentile of pMTnet-omni’s predicted rank% for TCR-pMHC pairs, and Gini indices of the average clonal sizes of TCRs targeting each pMHC, for each week in the subcutaneous tissue samples, lung tissue samples, and their averages. (e) Circos plots visualizing the pMTnet-omni-predicted TCR-pMHC binding pairs at week 6 and week 8 in the lung tissue samples. TCR clonotypes were visualized as orange pies, with the sizes of the pies referring to clonal expansion sizes. The pMHC were visualized as black pies, with the sizes of the pies referring to the number of unique TCR clonotypes that were predicted to be binding this pMHC. (f) Effector, exhaustion, and memory T cell marker expression and clonal expansion of T cells at subcutaneous and lung tissue samples for each week.

We then assessed the interactions between TCRs and tumor-associated antigens (TAAs). To define TAAs, we examined the expression of genes in tumor cells (both subcutaneous and lung) *vs.* all the other types of cells combined. We retained only those genes that are widely and highly expressed in tumor cells (>95% tumor cells, log2FC>3) and expressed in very few (<20%) normal cells. We further limit to genes that have been previously reported as putative tumor antigens in mice (**Sup. Table 4**). Using these criteria, we detected *Krt8* and *Krt18* as TAAs (**Fig. 2c**). We predicted the epitopes in *Krt8* and *Krt18* that bind to 129/Sv mouse MHCs by netMHCpan and netMHCpanII, and further predicted the mouse TCRs that bind *Krt8* and *Krt18* pMHCs, using pMTnet-omni.

We first calculated the proportions of TCRs predicted to bind *Krt8* and *Krt18* pMHCs. Consistent with a mounting anti-tumor immune response, in both the subcutaneous and lung sites, these proportions increased over time (**Fig. 2d**). We also examined the quality of the binding TCRs, reflected by rank%. For all TCR-*Krt8*/*Krt18* pMHC pairs from each tissue site and from each time point, we calculated the third percentile of all rank% scores (other percentiles yielded similar results). If there were no binding, the rank% would feature a uniform distribution and the third percentile of the rank% scores would be exactly 0.03. If there were an enrichment of binding pairs, the third percentile of rank% would be smaller than 0.03. As shown in **Fig. 2d**, the third percentile of rank% of all pairs were lower than 3% and decreased over time, reaching the lowest point around week 8. Furthermore, we investigated whether the TCR-*Krt8*/*Krt18* pMHC pairs demonstrated any immuno-dominance phenotype^31–33^ by using the Gini index to identify preferential expansion of TCR clonotypes towards certain pMHCs. A Gini index of 0 means perfect equality, while a Gini index of 1 means maximal inequality. As shown in **Fig. 2d**, we observed an increasing trend and a transient spike of Gini index at around week 5 in the subcutaneous site, which quickly settled in week 6. Then the Gini indices for both the subcutaneous and lung sites increased over weeks 7-9. We visualized the TCR-pMHC interactions (**Fig. 2e**), which were largely uniform at week 6 in the lungs. At week 8, however, the first and third most clonally expanded TCR clonotypes were both predicted to target the same pMHC. Notably, the pMHC targeted by these two TCRs, as well as the other pMHC targeted by the second most expanded TCR clonotype, were also the pMHCs attracting the greatest number of TCR interactions (namely, also counting non-expanding TCRs).

Immune cell receptor sequences correlate with the expression of the immune cells carrying these receptors^10,34^. Therefore, we examined the evolution of T cell gene expression with TCRs predicted to bind or not bind the *Krt8*/*Krt18* pMHCs. In **Fig. 2f**, we show that T cells in primary tumor sites generally have higher effector gene signatures than T cells in the lungs (marker gene lists provided in **Sup. Table 5**). The effector signature of binding T cells increased over time in primary sites, while it mostly stayed stable for non-binding T cells. In the lungs, the effector signatures of both binding and non-binding T cells increased. But after week 7 (*i.e.*, shortly after metastasis to the lungs), the binding T cells in the lungs started to demonstrate higher effector signatures than non-binding T cells. A similar dynamic was also observed for T cell exhaustion levels, with the peak of exhaustion signatures in binding T cells in lungs occurring around week 8. We also examined the clonal sizes of the TCR clonotypes. Interestingly, the clonal sizes of the binding T cells in the primary sites and the lungs both peaked around week 8 (**Fig. 2f**), and the clonal expansion is most dramatic in the primary sites.

Overall, our data suggest that the T cells in the subcutaneous sites and the lungs demonstrate evolution of clonal structures and changes in transcriptomic status in concert with tumor progression and initiation of metastasis. The largest change was observed around week 8, about 2-3 weeks after the emergence of metastasis in the lung. We also analyzed the interactions of TCRs with neoantigens in these mice, which are peptides derived from non-synonymous mutations presented by MHCs^35–37^ (**Sup. File 3 Fig. 2**). Here, pMTnet-omni indicated a similar time-course in the perturbation of the T cells and TCR repertoire ensuing metastasis.

### pMTnet-based Biomarker for Monitoring Risks of irAE of Cancer ICI Treatment

We next evaluated whether pMTnet can facilitate monitoring the evolution of other non-tumor-specific TCRs in cancer patients. To achieve this, we leveraged the predictive power brought by pMTnet-omni to develop a biomarker for the prediction of immune-related adverse events (irAEs), autoimmune toxicities that occur when immune checkpoint inhibitors (ICIs) unleash unwanted cytotoxic T cell responses against healthy organs. irAEs convey substantial morbidity, incur considerable costs, and in some cases may preclude further use of ICI drugs^38^. As more toxic combination ICI regimens are approved and immunotherapy use expands to earlier-stage cancers and smaller, less experienced community sites, such biomarkers could help inform the selection of treatment and monitoring.

We assembled a cohort of 57 cancer patients treated by ICIs (anti-PD1/-PDL1/-CTLA4, mono-or dual-therapy) in UT Southwestern Medical Center (UTSW) (**Fig. 3a**). Median age was 66 years, 32% were women and 67% patients developed grade ≥2 irAE. Tumor responses were evaluated using Response Evaluation Criteria in Solid Tumors (RECIST) guidelines. Complete response (CR), partial response (PR), stable disease (SD), and progressive disease (PD) were observed in 2%, 37%, 56%, and 4% of patients, respectively. A total of 134 blood samples were collected and cryopreserved at pre-ICI baseline; 3, 6, 8 weeks; every 12 weeks; and at the time of irAE occurrence. Patient clinical characteristics are summarized in **Sup. Table 6**. For each blood sample, we performed TCR-sequencing and cytokine profiling.

**Fig. 3.**
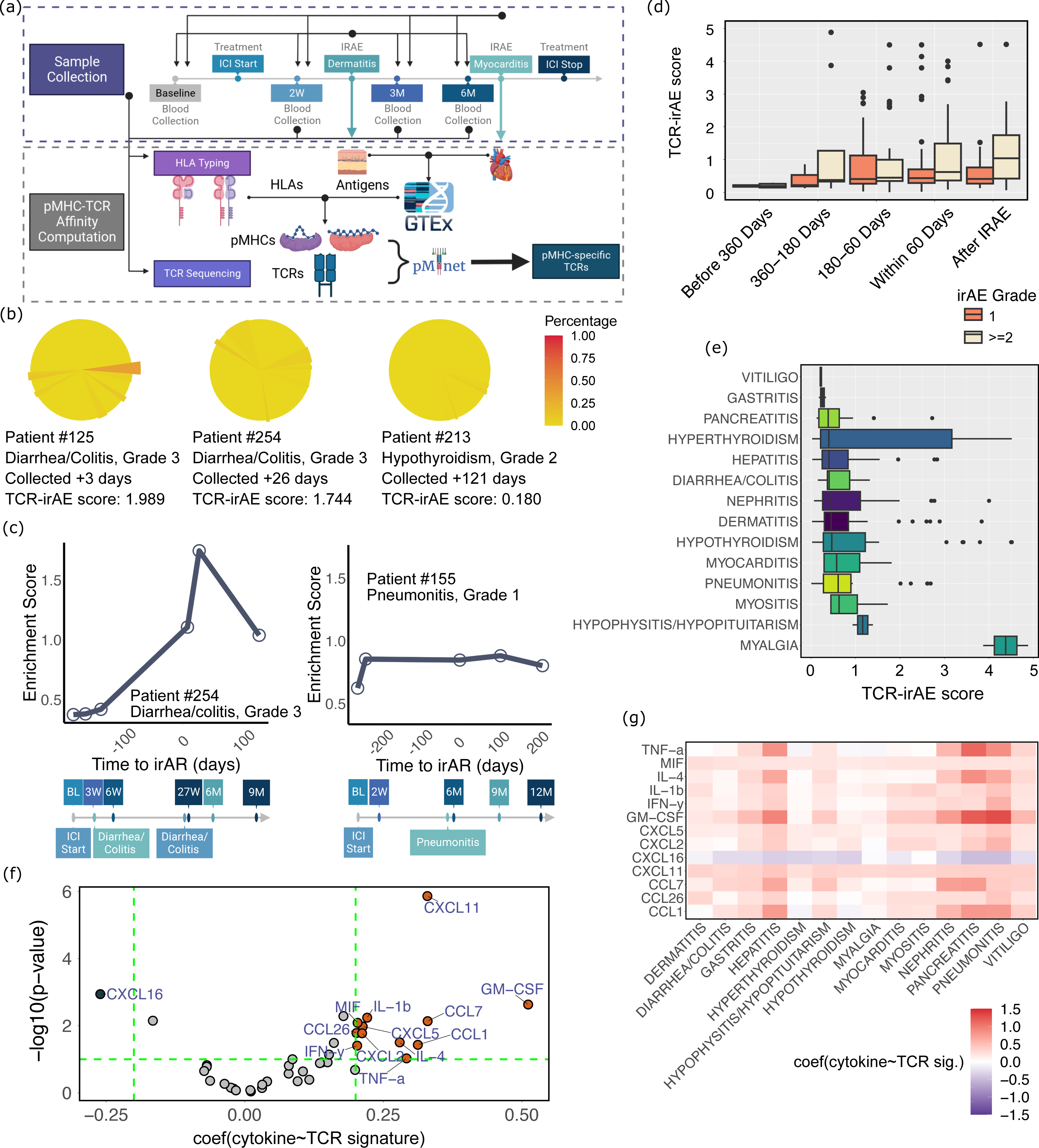
pMTnet-omni predicts irAE of ICI-treated patients. (a) Cartoon diagram showing the study design of how the patients’ blood was sampled and how data was generated. (b) Piecharts intuitively showing how the TCR-irAE prediction scores were calculated. Each TCR clone is shown as a pie, with redder color and greater radius meaning higher percentages of all pMHCs associated with the organ affected by the irAE are predicted as bound by this TCR. The clonal expansion sizes of the TCR clones are detonated by the fractions of the pies. (c) The TCR-irAE prediction scores overall blood sample collection time points of two patients. Time 0 in x-axis refers to the date of irAE occurrences. The timeline below shows the standardized sample collection time, ICI start time, and irAE occurance time. (d) The irAE prediction scores of all patients across all blood collection time points. The patient samples were separated into grade 1 and grade >=2. (e) The distribution of the TCR-irAE scores by irAE type. (f) A mix-effect model analysis testing how each cytokine’s level in the patients’ blood varies as a function of the TCR-irAE risk scores. Positive coefficient means a higher cytokine level when the TCR-irAE score is higher. (g) the mix-effect model’s coefficients specific for each irAE and each cytokine.

We first defined auto-antigens of healthy organs from the GTeX data^39^, which generated gene expression profiling for a total of 54 healthy tissues/organs. We defined putative auto-antigens as proteins that (a) are translated from genes expressed at least 10 times higher in one organ than the average expression of the same genes in all the other organs, and (b) have expressions of at least 50 TPM. For each putative auto-antigen, we used netMHCpan/netMHCIIpan to predict the peptides from these proteins that are presented by each patient’s specific MHCs. For each patient, pMTnet-omni was used to predict binding between the patient’s TCRs and the auto-antigenic pMHCs (**Fig. 3a**). We then defined a metric to measure whether patients’ top expanded TCRs are enriched with TCRs cytotoxic for each organ’s auto-antigenic pMHCs. To do so, we considered pMTnet-omni predictions and TCR clonal sizes. Details of this metric are described in the **method section**. **Fig. 3b** provides an intuitive visualization of this metric.

We then tested this approach in the two patients from our cohort with the greatest numbers of blood samples. In both cases, irAE risk scores increased over time and peaked around the time of irAEs diagnosis, especially for the patient with the higher grade (grade 3) irAE (**Fig. 3c**). When we aggregated all patients and samples from this cohort, we again observed the same trend of irAE scores increasing up to the time of irAE (Pval=0.01, one-sided Jonckheere’s trend test) (**Fig. 3d**). When we segregated patients according to mild irAE (grade=1) and those with more severe irAE (grade>=2), we observed higher irAE risk scores in cases with higher-grade irAE (Pval=0.013 for samples from all time windows combined, T-test) (**Fig. 3d**). These results suggest that the pMTnet-omni-based risk score is indeed predictive of irAE. Across irAE types, risk scores were highest for myalgia, hypophysitis/hypopituitarism, myositis, pneumonitis, and myocarditis, suggesting that these irAEs may be the most T cell/TCR-dependent (**Fig. 3e**). Interestingly, myocarditis has been associated with epitope-specific T cell responses in mouse models of myocarditis irAE^40^.

To investigate the relevance of the irAE scores computed based on TCR sequences for T cell functions, we explored the patient cytokine profiles. We correlated risk scores with cytokine levels by regressing each measured cytokine on computed irAE risk scores. By examining the resulting coefficients and corresponding P-values, we observed that higher irAE scores were positively associated with up-regulation of pro-inflammatory cytokines, including CXCL11^41^, GM-CSF^42^, CCL7^43^, IL-1b^44^, *etc* (**Fig. 3f**). Namely, our results reassuringly confirm that the patients’ immune systems are in an inflammatory state when the irAE risks are high, again corroborating our approach. Furthermore, we visualized the correlations for each specific irAE category, and found that CXCL11 and MIF which are also generally regarded as pro-inflammatory^45^, were the most consistently associated with variations of irAE risk scores across irAE categories (**Fig. 3g**).

### Preferential Mobilization of Prior SARS-CoV-2-specific TCRs after COVID-19 Vaccination

To further validate the potential use of pMTnet-omni to monitor the functional relevance of the evolving TCR repertoire in various contexts, we analyzed patients after COVID-19 vaccination. Specifically, we studied a set of 7 patients who recovered from prior SARS-CoV-2 infections and later received mRNA-based COVID-19 vaccines^46^ (**Fig. 4a**). Vaccines were administered an average of 434 days after onset of symptoms related to their prior SARS-CoV-2 infection. Blood samples were taken prior to and post vaccinations, with blood collection times shown in **Fig. 4a**. Multiplex scRNA-seq/scTCR-seq was performed on the sorted T cells. We predicted binding of viral epitope proteins (spike, replicase, nucleoprotein, membrane, envelope, and other ORFs) by patient-specific MHCs, and further predicted binding of presented pMHCs by each patient’s TCRs with pMTnet-omni. First, we assessed whether spike peptide-specific TCRs have higher clonal expansion (**Fig. 4b**, mixing all patient samples and mixing CD8^+^/CD4^+^ T cells). Strong binding TCRs were significantly more clonally expanded than weak and non-binding TCRs (Pval=0.03 and 0.01, respectively, one-sided Wilcoxon test). Given this, we further evaluated strong binding TCRs in different T cell subsets.

**Fig. 4.**
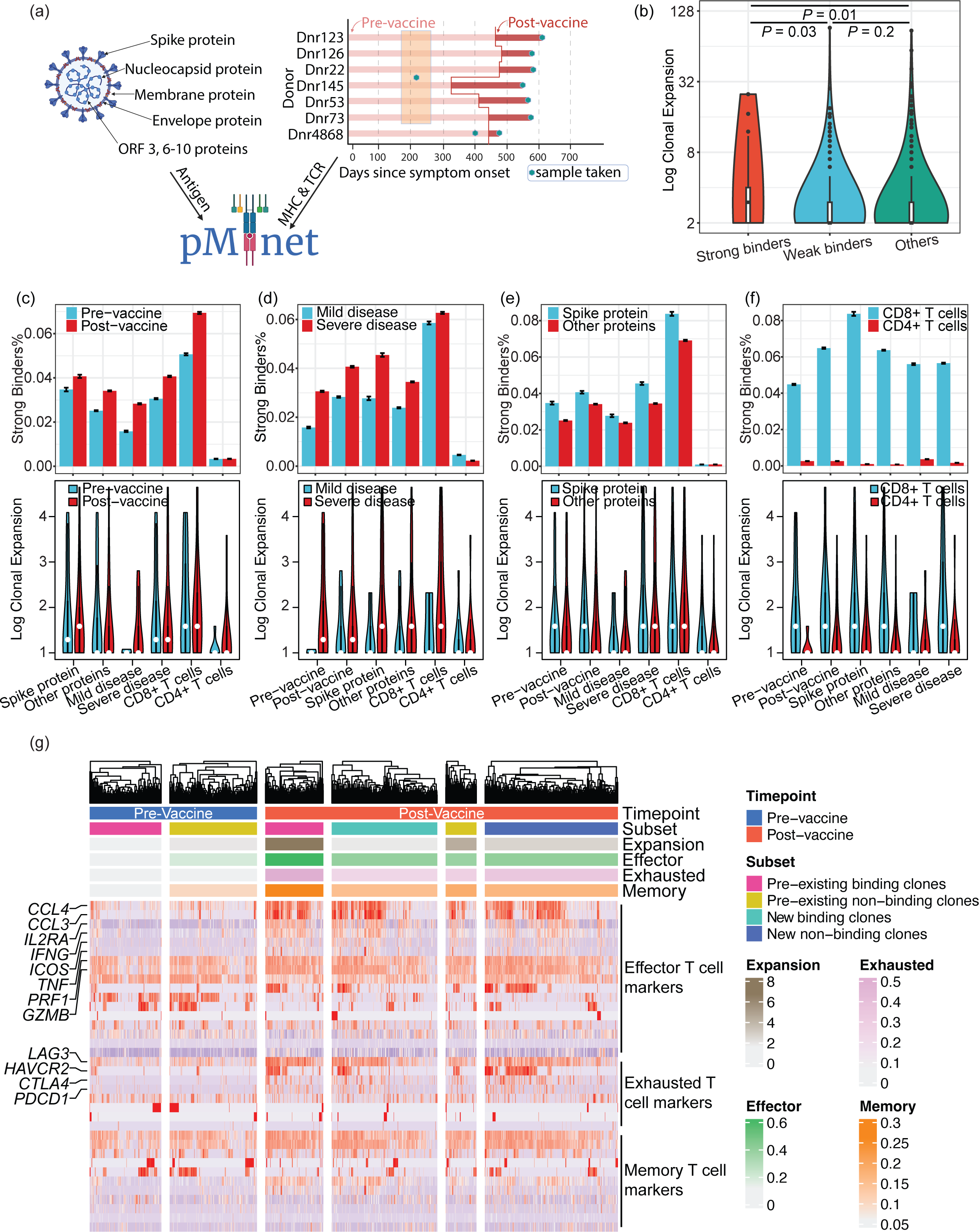
pMTnet-omni interprets TCR clonal evolution in patients with SARS-CoV-2 infection and vaccination. (a) Cartoon diagram showing the study design. (b) Clonal expansion of T cells predicted to target SARS-CoV-2 spike protein. Binding T cells were classified into strong binders (rank%<=0.05%) and weak binders (rank%<=3%), and non-binding T cells (rank%>3%) were also shown as a negative control. P values were calculated by one-sided Wilcoxon test. (c) The impact of vaccination on binding affinity (percentage of strong binders) and clonal expansion of T cells. (d) Binding affinity and clonal expansion of T cells in patients with mild or severe COVID-19 disease. (e) Binding affinity and clonal expansion of T cells targeting spike protein and other proteins of SARS-CoV-2. (f) Binding affinity and clonal expansion of CD8^+^ and CD4^+^ T cells. In panels c-f, data from patient Dnr4868 were removed where the comparison of mild *vs* severe disease is concerned, as this patient was not designated as having severe or mild prior SARS-CoV-2 infections in the original data. (g) Effector, exhaustion, and memory T cell marker expression and clonal expansion of 4 subsets of CD8^+^ T cells pre- and post-vaccination. T cell clones appearing in the pre-vaccination samples were grouped into SARS-CoV-2-specific (pre-existing binding clones) and non-binding (pre-existing non-binding clones) subsets. T cell clones appearing in the post-vaccination samples but not in the pre-vaccination samples, were grouped into SARS-CoV-2-specific (new binding clones) and non-binding (new non-binding clones) subsets.

We subset strong binding TCRs into those from T cells of pre-vaccination samples and of post-vaccination samples (**Fig. 4c-f**). As expected, TCRs from post-vaccination samples were more likely to bind to SARS-CoV-2 viral epitopes than TCRs from prior-vaccination samples and they were also more clonally expanded. We compared patients with mild (non-hospitalized) *vs.* severe (hospitalized) prior SARS-CoV-2 infections. T cells from patients with severe infections were more likely to be viral epitope-binding and more clonally expanded. We also analyzed patterns of CD8^+^ T cells and CD4^+^ T cells. As is shown in **Fig. 4c-f**, CD8^+^ T cells were the dominant population responding to SARS-CoV-2. Next, we investigated pMHCs from SARS-CoV-2 spike and other viral proteins. We found that T cells respond to both spike and other viral proteins, but have a preferential response to spike proteins (**Fig. 4c-f**).

We also compared T cells and TCR clonotypes pre- and post-vaccination. We divided the SARS-CoV-2 protein-binding or non-binding TCRs into TCR clonotypes present in pre-vaccination samples and those that emerged only after vaccination. We first examined the clonal expansion of these TCR clonotypes. **Fig. 4g** shows that pre-existing SARS-CoV-2-specific TCR clones (maroon subset) were highly expanded after vaccination, while much less expansion is observed for pre-existing TCR clones that are not SARS-CoV-2-specific (yellow subset). Notably, newly appearing binding TCR clones (bluey-green subset) were not as expanded as pre-existing binding clones. Concomitant with clonal expansion, T cells with pre-existing binding clones had stronger expression of activation, exhaustion, and memory markers after vaccination, compared with pre-vaccination (marker gene lists provided in **Sup. Table 5**). Again, in post-vaccination T cells, pre-existing binding T cells also had higher activation/exhaustion/memory phenotypes than the T cells with newly appearing binding TCR clones (**Fig. 4g**). Taken together, these results suggest that the human immune system preferentially mobilizes pre-existing SARS-CoV-2 targeting T cells, rather than newly emerging SARS-CoV-2-specific T cells, in these vaccinated patients.

### Delineating the Spatial Heterogeneity of TCR-antigen Interactions

While our previous analyses focus on longitudinal tracking of TCRs, we next used pMTnet-omni to characterize spatial heterogeneity of TCR-antigen interactions from spatial-TCR-sequencing data. We analyzed a Visium-TCR-seq dataset from Hudson *et al*^47^. Visium-TCR-seq is a variation based on 10X Visium technology that provides information on gene expression, spatial location, and TCR clonotypes for thousands of sequencing spots. This Visium-TCR-seq dataset is derived from the brain metastasis of a melanoma patient, and contains a total of 1,716 sequencing spots with expression data on 36,601 genes and 140 unique TCR clonotypes, as well as matching H&E staining image (**Fig. 5a**). We first performed cell typing for this Visium-TCR-seq dataset. As each Visium sequencing spot contains ∼10 cells of potentially different types, it is difficult to assign cell types to the highest resolution, as can be seen from the UMAP plot of gene expression (**Fig. 5b**). However, we were able to reliably identify tumor, stromal/immune, and blood cell sequencing spots, based on gene expression and pathological appearance.

**Fig. 5.**
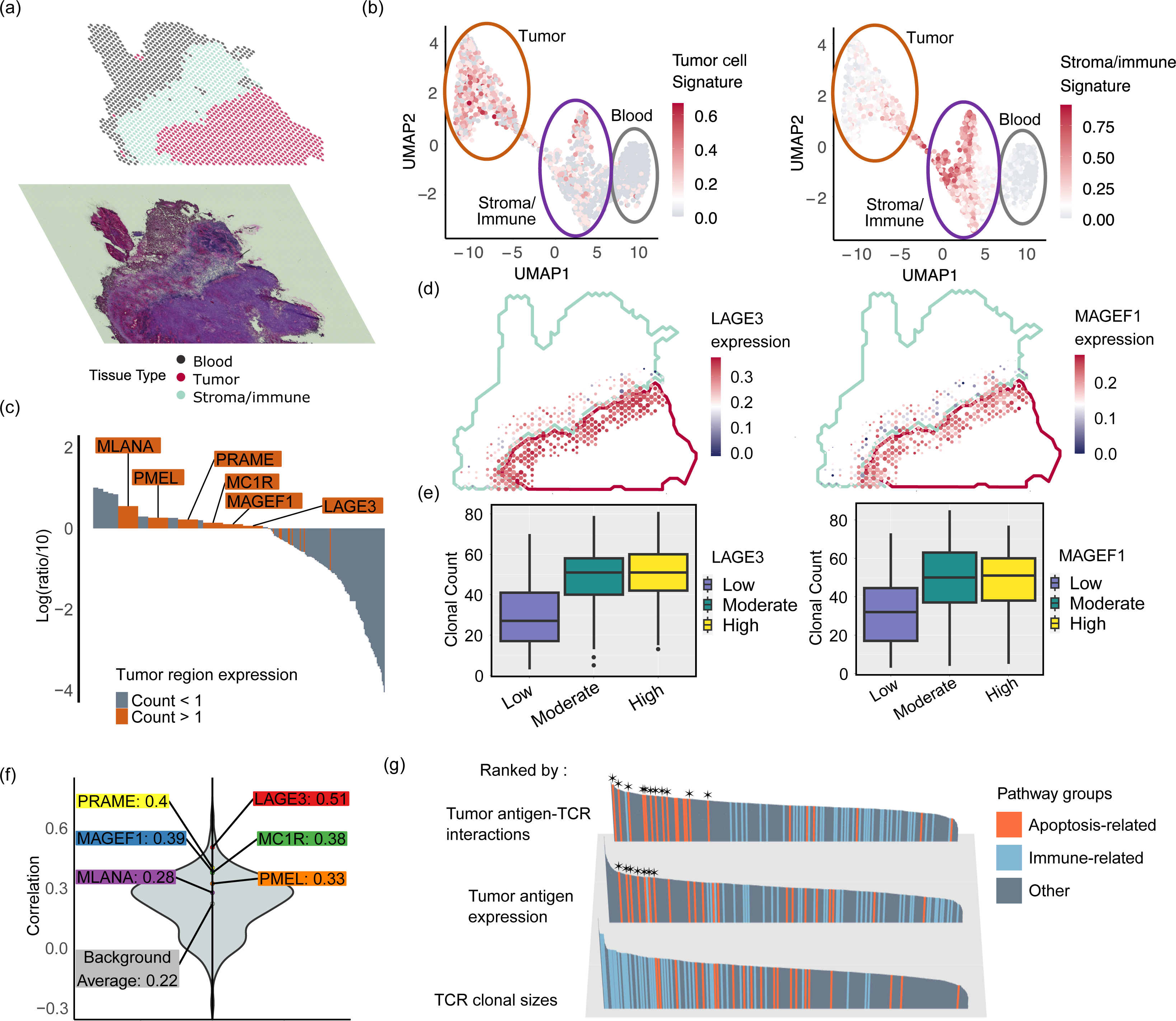
pMTnet-omni characterizes the intra-tumor spatial heterogeneity of TCR-pMHC interactions. (a) The Visium-TCR-seq dataset, showing the spatial spots with gene expression information and the corresponding H&E staining image. The spots were divided into tumor, stroma/immune, and blood spots based on gene expression. (b) Umap clustering of the Visium sequencing spots, with the color denoting expression of the tumor and stroma/immune marker genes, respectively in the two figures. (c) Genes with tumor/non-tumor expression fold changes greater than 10, average expression counts in the tumor spots greater than 1, and previously described as TAAs were retained for the following analyses. (d) Visualization of the spatial distribution of the Visium-sequencing spots near the tumor-stroma/immune interface, with dot sizes referring to TAA-specific TCR clonal sizes and color denoting TAA expression. (e) Boxplots showing the clonal sizes of TAA-specific TCR clonotypes in tumor sequencing spots near the tumor-stroma/immune interface, with these sequencing spots divided into three categories based on the level of TAA expression. (f) Correlations between TCR clonal sizes and TAA expressions of the tumor sequencing spots near the tumor-stroma/immune interface. (g) Gene Set Enrichment Analysis (GSEA) to infer pathways that are enriched in the genes that are differentially expressed in Visium sequencing spots near the tumor-stroma/immune interface, between the sequencing spots of high *vs* low TAA-TCR interactions, with high *vs* low TAA expression or with high *vs* low TCR clonal expansion. Pathways that are related to apoptosis with p-values <= 0.05 are marked by asterisks.

Next, we investigated interactions between tumor-associated antigens (TAAs) and T cell receptors. We first defined TAAs by examining genes highly expressed in tumor regions as opposed to stroma/immune/blood regions (**Fig. 5c**). We required a minimum average count of 1 in tumor regions and a minimum tumor-stroma/immune/blood expression ratio of 10. We also limited our search to genes previously described as putative TAAs (**Sup. Table 3**). Six genes satisfied all these criteria: *MLANA*, *PMEL*, *PRAME*, *MC1R*, *MAGEF1*, and *LAGE3*. We predicted the pMHCs from these TAAs by using netMHCpan and netMHCIIpan, with HLA alleles typed by arcasHLA^48^. We then predicted interactions between these TAA pMHCs and TCRs using pMTnet-omni.

While we showed above that TAA-specific TCR clones arise during tumor progression, the Visium-TCR-seq here allows us to determine whether TAA-specific T cells actively co-localize with tumor cells that highly express TAAs. We examined the expression of the TAAs, and the clonal expansion of TAA-specific TCR clonotypes, for the sequencing spots near the tumor-stroma/immune interface, where T cells and tumor cells predominantly interact (**Fig. 5d**, details of calculation in **method section**). We discovered that higher TAA expression is associated with higher TCR clonal expansion in these sequencing spots (**Fig. 5e**). We quantified these correlations for each of the six TAAs and calculated a background distribution by random mismatching the TAA-specific TCRs with 300 other genes captured in the dataset (**Fig. 5f**). We confirmed that correlations between TAAs and TAA-specific TCRs are all higher than the average correlation of the background distribution (Pval=0.002, one-sided T test). Therefore, pMTnet-omni revealed the active co-localization of TAA-specific TCRs with TAA-expressing tumor cells. We also analyzed a Slide-TCR-seq dataset from Liu *et al*^49^, and made similar observations (**Sup. File 3**).

The expression component of the spatial-TCR-seq data also allowed us to examine whether the predicted TAA-TCR interactions correlate with transcriptomic changes. For the Visium sequencing spots highlighted in **Fig. 5d**, we calculated a TAA-TCR interaction score based on the numeric product of TAA gene expression and the clone sizes of TAA-specific TCRs, and computed the average score across all six TAAs. We divided these sequencing spots into high-interaction and low-interaction spots. As a control, we computed a TCR clonal expansion score, by considering only the average clonal sizes of all TCRs in each sequencing spot, and similarly divided all sequencing spots into high-TCR and low-TCR spots. We also created another control, by dichotomization based on the sum of expression of TAAs in the sequencing spots. Gene Set Enrichment Analysis (GSEA)^50^ was performed to compare the gene expression between the two subsets of spots and to compute enriched pathways. Interestingly, we observed that many top pathways enriched in the analysis comparing high *vs.* low TAA-TCR interactions were related to cell death, particularly apoptosis (**Fig. 5g**, the apoptosis-related pathways with GSEA Pval<=0.05 are marked by asterisks), which is reflective of the cytotoxic functions of the T cells. In comparison, top pathways enriched in the analysis comparing high *vs.* low TCR clonal sizes were only related to immune responses. Cell death-related pathways were also less enriched in the comparison of high *vs* low tumor antigen expression (Pval=0.018, Kolmogorov-Smirnov test), as opposed to the comparison of high *vs* low interactions (Pval=0.00033). These results indicate that T cells with predicted TAA-specific TCRs were competent in their cytotoxic functions and induced tumor cell death.

### pMTnet-omni Recognizes Small Differences in TCR Sequences

While we have demonstrated that pMTnet-omni can determine whether a TCR binds or does not bind a pMHC, we further explored whether pMTnet-omni can recognize small differences in TCR sequences, which likely result in smaller shifts in binding affinity. We examined the four CD8^+^ T cell scRNA/scTCR datasets from 10X, which contain extra data on binding affinity of each TCR toward 44 class I pMHCs. The binding affinity information is captured by 10X’s feature-barcoding technology and is recorded by a Unique Molecular Identifier (UMI) count, with higher count denoting potentially stronger binding. To validate the functional relevance of UMI counts, we confirmed that a higher UMI binding score is indeed correlated with stronger clonal expansion (**Fig. 6a**). Gene Ontology (GO) analyses revealed that genes differentially expressed between T cells with binding *vs.* non-binding TCRs, defined by UMI counts, are enriched in pathways related to T cell activation (**Fig. 6b**). And the binding T cells indeed have higher expression of T cell activation marker genes (**Fig. 6c**).

**Fig. 6.**
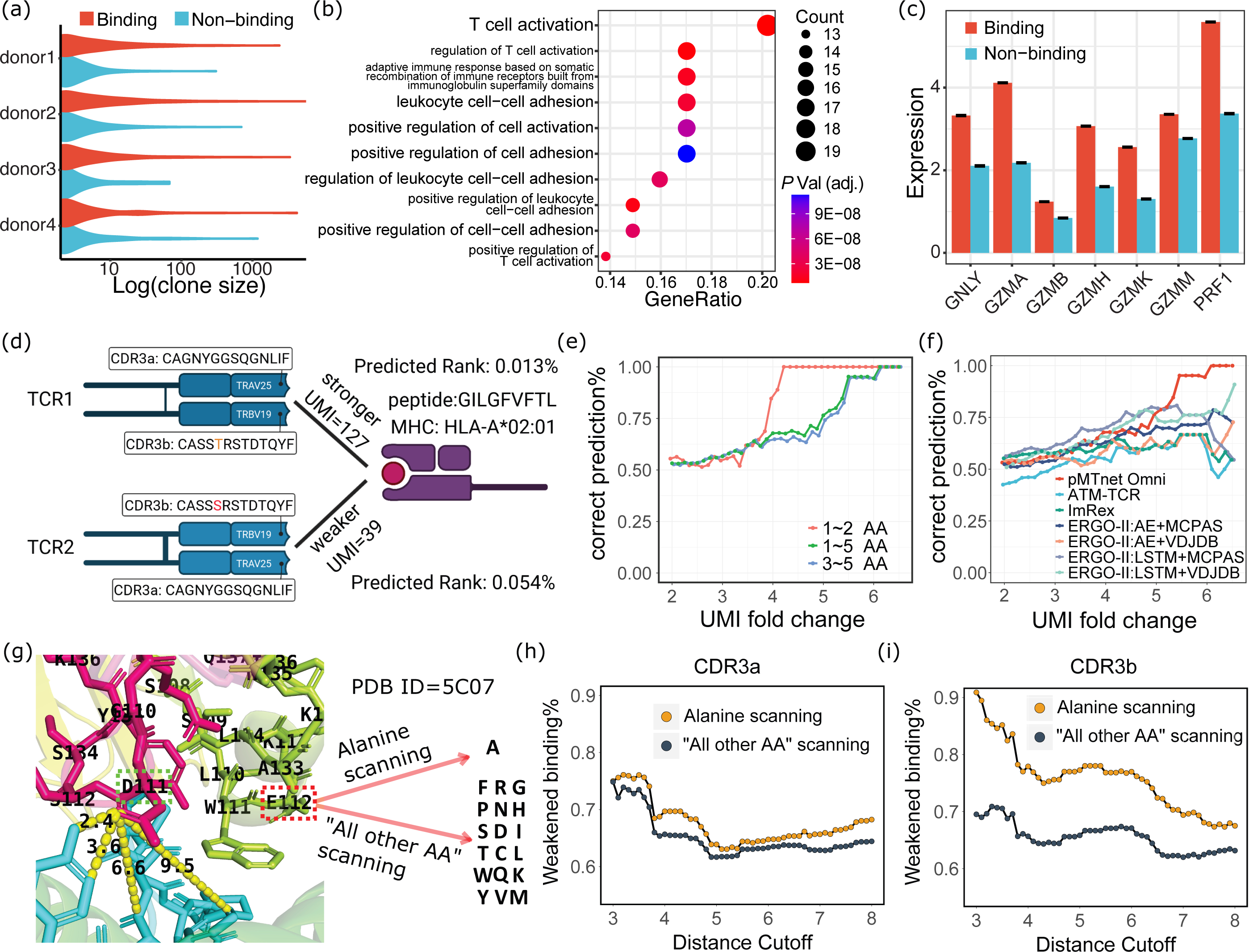
pMTnet-omni distinguishes the binding affinity of similar TCRs. (a) Clonal expansion of binding and non-binding TCR clones in each patient. (b) Gene Ontology analyses of pathways enriched in the differentially expressed genes between the binding and non-binding T cells. (c) Expression of T cell activation marker genes and cytotoxicity-related genes in binding and non-binding T cells. (d) Diagram showing an example pair of TCRs sharing similar sequences but with different binding affinity measured by UMIs. (e) Percentage of correct prediction for the similar TCR pairs. The X-axis represents the minimum fold change of the better binding TCR over the worse binding TCR in terms of UMI counts. (f) Benchmarking correct prediction rates for the similar TCR pairs for several competing software applications. (g) 3-dimensional structure of an example TCR-pMHC pair (PDB ID=5C07), demonstrating how the distances between CDR3 residues and pMHC residues were calculated, for D111, and showing how “Alanine scanning” and “All other AA scanning” were performed, for E112. (h) Line plots showing the percentages of predicted weakened binding for the TCR-pMHC pairs, as a function of distance between the CDR3 residues and pMHC residues.

Next, we extracted pairs of similar TCRs that bind the same pMHCs from this dataset, where the two TCRs in each pair are both binding, share the same V genes and differ by no more than 5 amino acids (AAs) in their CDR3α and CDR3β sequences combined (**Fig. 6d**). The two TCRs in each pair were designated as stronger and weaker binders, according to the feature-barcoding UMI counts. We then assessed whether pMTnet-omni correctly identified the stronger/weaker binders. As shown in **Fig. 6e**, the prediction accuracy for discerning stronger/weaker binders increases as a function of the differences in binding affinity between the two TCRs, measured by UMI counts. Namely, as two TCRs demonstrated larger differences in pMHC binding (considering that this UMI count likely is not perfect and has noises), the probability of correct prediction by pMTnet-omni was also higher. We segregated all TCR pairs into those with 1-2 AA differences and those with 3-5 AA differences. For both subsets, we observed the same increase in prediction accuracy (**Fig. 6e**). We also executed the benchmark software from **Fig. 1f** and compared their performance against pMTnet-omni. As **Fig. 6f** indicates, pMTnet-omni had above-average performances compared with the benchmark software when UMI fold change cutoffs were lower, and out-performed all benchmark software by a large margin when only considering TCR pairs with higher UMI fold change cutoffs. When subsetting the TCR pairs into those with 1-2 AA and those with 3-5 AA changes, we observed similar patterns (**Sup. Fig. 7**). Interestingly, the prediction accuracy for TCR pairs with 1-2 AA differences was higher than those with 3-5 AA differences at the same level of UMI fold change. The AA changes in the TCR pairs with 1-2 AA differences likely were positioned on more important residues than those TCR pairs with 3-5 AA differences but achieving the same level of UMI fold change (the latter subset of TCRs need more AA changes to achieve the same level of UMI differences).

### Therefore, pMTnet-omni might have attended more to the more important TCR residues

To corroborate whether pMTnet-omni can truly distinguish residues according to their importance, we studied 155 three-dimensional structures of TCR-pMHC complexes from TCR3d^51^. We performed an *in silico* “alanine scanning” (**Fig. 6g**), a common approach taken in biophysics studies^52,53^, by switching each AA on the CDR3s to alanine. Alanine has a small side chain and is chemically inert. Therefore, if the original AA is important for binding, switching to alanine will likely impact binding negatively. We then employed pMTnet-omni to compute the binding strength of the mutated TCRs. As shown in **Fig. 6hi**, most mutated TCRs yielded weaker predicted binding strength, consistent with our hypothesis. Notably, the probability of observing diminished binding was stronger when the ablated CDR residues were physically closer to pMHCs residues (X axis of **Fig. 6hi**, unit: Ångströms), and the probability decreased over increasing distances. This is because the residues on TCRs that are spatially closer to pMHC residues are more likely to be the key residues impacting binding. Moreover, by comparing the alanine scanning results for CDR3α (**Fig. 6h**) and CDR3β (**Fig. 6i**), we observed that alanine scanning of CDR3β residues, which are more important for binding (**Fig. 1e**), is more likely to result in diminished binding, again confirming our conclusion. We also performed “other AA scanning”, by switching each CDR3 AA to other non-alanine AAs. As **Fig. 6hi** indicates, the same conclusions can be drawn, though with less magnitude of differences.

## DISCUSSION

In this work, we developed pMTnet-omni with state-of-the-art pan-MHC and cross-species TCR-pMHC pairing prediction performance, benefitting from the innovative model design and training procedures, as well as careful curation of the pairing data generated by the collective efforts of all researchers in the field. pMTnet-omni can distinguish binding *vs.* non-binding TCRs against a pMHC, and can also distinguish stronger binding *vs.* weaker binding TCRs. Furthermore, we demonstrated the potential of developing diagnostic biomarkers by building upon pMTnet-omni, for longitudinal and functional monitoring of the TCR repertoire during disease progression and therapeutic interventions. While our study presented here focuses on ICI-associated irAE, COVID-19 vaccination, and metastatic melanoma, we believe the same principle can be extrapolated to other clinical scenarios in which T cells play TCR-dependent roles.

Fast and accurate computational prediction of TCR-antigen binding will accelerate progress toward real-time monitoring of the antigen binding landscape of individual TCR repertoires and generation of personalized TCR-based therapeutics. This is especially important, given that TCRs bind antigens with relatively lower affinity and lower specificity^54–57^, compared with binding of BCRs/antibodies towards antigens. We must view TCR-antigen pairing relationships as networks, where multiple TCRs can target one antigen and multiple antigens can be targeted by the same TCR. This necessitates the capability to map TCR binding of antigens *via* some machine learning-based systems, such as pMTnet-omni, as experimental assessment of all possible TCR-antigen pairs is prohibitive cost- and time-wise, or may not even be feasible logistically.

pMTnet-omni employed a unique hybrid sequence-structure framework to solve the prediction of TCR-pMHC pairings. This approach enabled us to consider the structural features of the binding complexes, but without being limited by the very small number of solved TCR-pMHC protein structures. We modeled MHCs through a structure submodel, and TCRs and peptides through sequence submodels, based on our understanding of the degree of protein disorderedness in the local regions where TCRs (CDRs) and pMHCs interact. Nevertheless, this is not the only reasonable design, and we will explore alternative approaches in the future. For instance, CDR and peptide conformations are likely to become more rigid upon binding of TCR-pMHCs and packing of the amino acid residues in the limited space. Therefore, structural submodels might also be suitable for modeling CDRs and pMHCs, or could at least provide some information that has not been captured in sequence-based models. We imagine that the field could generate various models of TCR-pMHC pairing that are purely sequence-based, purely structure-based, or that contain a mixture of structure and sequence information to various degrees. Then a “wisdom of the crowds”^58–60^ method can be taken to ensemble these complementary models, which will likely yield the optimal super-model for TCR-pMHC binding prediction.

TCR-pMHC pairing prediction models, such as pMTnet-omni, could be used to generate TCR immunotherapeutics. Despite being imperfect so far, these “dry” approaches offer advantages that cannot be provided by traditional “wet” methods. Take TCR-T therapies or TCR-like drugs for example. Most TCR-T therapies simply use TCRs isolated from patient blood that target the antigen of interest. This approach is taken because TCR optimization against target antigen— while possible—is difficult, time-consuming, and expensive. Even if one changes only *N* (*N*=*1, 2, 3* for example) amino acid residues in the CDR3s of the α and β TCR chains, which total an average of ∼50 amino acids, the search space will be around ((19+20+1)x50)^N^, given 19 possible choices of amino acid substitutions, 20 choices of insertions and 1 choice of deletion. And this is just one round of optimization against one target antigen. Further, there are many possible off-target auto-antigens in the human body that might be targeted by the variant TCR. To increase affinity against the target antigen and decrease affinity against off-target antigens, the total amount of screening efforts multiplies. Further, future TCR engineering efforts might want to consider modification of TCRs so that they are more compatible with the HLA alleles of more patients, or generate a set of TCRs that collectively target one antigen. At that point, screening tasks become magnitudes more difficult or even impossible for traditional “wet” lab experiments. However, such tasks can be accomplished by simply running AI prediction programs longer and coding desired TCR properties into a loss function against which the AI model is optimized. Even if such prediction models are not perfect, they will benefit the TCR discovery efforts as long as they can generate a small list of prioritized TCRs that can be subsequently validated with a reasonable validation rate.

Nevertheless, neither pMTnet-omni nor similar works completely replace the need for wet lab validations. Rather, still considering TCR-T therapies or TCR-like drugs, one might envision an iterative process between AI prediction and experimental validation to be the most productive route. With the aid of AI, the scale of the validation experiments can be much smaller. The validation outcome, either positive or negative, can be fed into machine learning models for fine-tuning, which will generate more accurate predictions for the next iteration. Furthermore, some non-TCR factors could also influence binding of TCRs against pMHCs, such as TCR expression/clustering on cell surface and expression/function of co-stimulatory molecules, *etc*^57,61^. In other words, TCR-pMHC binding rules could vary, to some extent, across different biological conditions, such as different cultures or stimulation conditions. Unfortunately, it is challenging for TCR-pMHC binding prediction models to consider these highly case-specific variables. Nevertheless, limited experiments in these biological conditions could generate small-scale TCR-pMHC pairing data that can be used to fine-tune an AI agent from the general pMTnet-omni model, to adapt to the specific condition under investigation and to generate successful predictions.

Finally, our work on TCR-pMHC pairing adds to the literature on prediction of protein/ligand binding. While works such as AlphaFold2 provide functionalities to solve more general protein/ligand binding prediction problems, they perform less well under specialized situations, such as those involving T and B cell receptors^62^. This is because these protein structure prediction models mostly rely on multiple-sequence-alignment and evolutionary modeling. However, T/B cell receptors are diverse and “somatic”, which are created through VDJ recombinations and somatic hypermutations, and the binding epitopes are also highly diverse. Due to these hurdles, dedicated works, such as pMTnet-omni, are called for to address these grand scientific questions and to generate immunotherapeutic products or diagnostic tools.

## METHODS

### Training and validation of the variational autoencoders for V genes and CDR3s

V gene alleles and amino acid sequences used for training V gene encoders were collected from the IMGT database. CDR3 amino acid sequences used for training CDR3 encoders were collected from the data sources listed in **Sup. Table 1**. VAE and denoising VAE models were employed to generate CDR3 and V gene embeddings, respectively. Due to the very few numbers of V gene alleles, data augmentation was used to increase the data size of V gene sequences. Amino acid substitution, insertion, and deletion were added to the original V gene allele sequences. After that, the augmented V gene allele sequences/CDR3 sequences were converted to Atchley Factor matrices and input to the VAE models. The VAE encoder for V gene consisted of a transformer layer and 10 convolution blocks. The VAE encoder for CDR3 consisted of a transformer layer and 25 convolution blocks. The bottleneck layers of both VAE models were extracted as the embeddings of V genes/CDR3 sequences. The reconstructed Atchley Factor matrices were compared with the Atchley Factor matrices of the original sequences to calculate the reconstruction loss. The total loss was the weighted sum of reconstruction (mean squared error, MSE) loss and Kullback–Leibler loss. During the validation process, the embeddings were computed only from the mean of the latent space (without the variational part of the VAE). The training and validation data for the V gene models were the V gene allele sequences retrieved from IMGT^63^. The training and validation data for the CDR3 models were CDR3 sequences collected from multiple sources listed in **Sup. Table 1**. There was no overlap between the training and validation sequences. Description of extra model architecture and parameter details could be found in **Sup. File 1**.

### Training and validation of the pMHC embedding model

The protein sequences of the α MHC subunit and the β subunit are input into the ESM2 650M model^20^ for numeric embeddings, in the case of class II MHCs. For class I MHCs, the MHC protein sequence and microglobulin protein sequences were used for ESM2 embedding. The peptide sequences were converted to Atchley Factor matrices. The MHC representations and Atchley Factor matrices of the peptides were input to the pMHC embedding model, which consisted of 9 convolution blocks. The head layer and the loss function of the pMHC model are constructed to predict the binding between the peptides and the MHCs. Then the embeddings of the pMHC complexes were extracted from the last layer before the prediction head layer of this model. The MSE loss and cross-entropy loss were used for the binding affinity (BA) training data and the eluted ligands (EL) training data, respectively. Description of extra model architecture and parameter details could be found in **Sup. File 1**.

The p-MHC binding model was evaluated with the validation dataset that was used by the original publication of NetMHCpan for validation of binding between peptides and human class I MHCs (this dataset contains human pairs only), and also mouse class I p-MHC binding data downloaded from IEDB. For class II MHCs, only p-MHC binding data downloaded from IEDB were used for evaluation because nearly all (916 out of 917) evaluation data points in the validation dataset from the netMHCIIpan publication were human HLA-DR alleles. The IEDB dataset was filtered to include class I and class II MHC-presented peptides with 8-30 and 9-30 amino acids, respectively. Peptides that comprised only the 20 common amino acids were retained. p-MHC pairing data that appeared in the training set were excluded from the evaluation set. Finally, there were 609,106 p-MHC pairs in the IEDB human class II MHC dataset, and 58,175 p-MHC pairs in the IEDB mouse class I/II MHC datasets.

### Training and validation of the pMHC-TCR binding prediction model

The VAE embeddings of Vα, Vβ, CDR3α, and CDR3β were concatenated and projected to a latent space by fully connected layers. The pMHC embeddings were also projected to latent space. Embeddings in the latent space were normalized to unit hypersphere, where dot product was used to measure the cosine distance between the TCR and pMHC numeric representations in the latent space. Supervised contrastive loss was used to train the pMHC-TCR model. The loss function was constructed such that the cosine distance between the positive (binding) TCR and this pMHC was expected to be smaller than the cosine distances between randomly sampled TCRs (negative TCRs) and this same pMHC. For 1 pMHC complex, 50 randomly sampled TCRs were contrasted with 1 truly binding TCR. The training process was repeated 3 times with 3 different random seeds. After training, these 3 pMHC-TCR models were ensembled by averaging the cosine similarities, divided by a scalar temperature parameter. During the validation process, a large number of species-specific TCRs were randomly sampled as the background TCRs for 1 query pMHC-TCR pair. The predicted binding score of the query pMHC-TCR was compared with the binding scores predicted for the binding between the query pMHC and all the background TCRs, to generate a rank% for the query TCR-pMHC pair. Smaller rank% denotes stronger predicted binding affinity. Description of extra model architecture and parameter details could be found in **Sup. File 1**.

### Generation and analysis of scRNA/scTCR-seq data from the 344SQ/129Sv mouse model

All animal experiments were approved by the Institutional Animal Care and Use Committee at MD Anderson Cancer Center. Wild-type 129Sv mice aged 7-8 weeks were subcutaneously injected with the 344SQ cell line, a mouse lung adenocarcinoma cell line originated from subcutaneous metastasis of genetic mouse models with *Kras*^G12D/+^;*p53*^R172HDG^ alleles^29^. Specifically, 0.5×10^6^ cells were administered in 100μL of phosphate-buffered saline (PBS) into the posterior flank^64^. Mice were regularly monitored and euthanized if they showed signs of morbidity or if the tumor diameter reached 15mm. Fresh samples were collected at intervals ranging from 3 to 9 weeks after inoculation. Specifically, mouse whole blood was drawn *via* intra-cardiac puncture into a K_2_EDTA vacutainer and then stored on ice. The mouse lungs were perfused using sterile PBS *via* heart perfusion from the left ventricle before the entire lung was harvested. Primary tumors from the mice were surgically excised and stored in RPMI-1640 awaiting further processing.

Solid tissues were finely minced into small pieces followed by using dissociation mixture containing 1mg/ml Collagenase A and 0.4 mg/ml Hyaluronidase in RPMI1640 with glutamine. After incubating on a rotary shaker at 37°C for 90 min, the mixture underwent further dissociation using 10ml of Trypsin-EDTA, and then 5ml of pre-warmed Dispase (5U/ml) and DNase I solution (50μl). The tissues were passed through to a 70mm cell strainer (Thermo Fisher Scientific) to obtain single-cell suspension. For the blood processing procedure, blood samples were combined with an equal volume of PBS and gently overlaid on the top of the Lympholyte Cell Separation Media (Lympholyte-M) and then centrifuged at 1,200 rcf for 20 min at room temperature. After centrifugation, a well-defined lymphocyte layer formed at the interface, carefully remove the cells using a pipette from the interface and transfer to a 50-ml tube. The cells were diluted with 20 ml PBS and centrifuged at 800 rcf for 10 min to pellet the lymphocytes.

The resulting cell isolates were treated with 10ml red blood cell (RBC) lysis buffer (BioLegend). The cells were resuspended in PBS with 1% BSA, adjusted to the density of 0.7-1.2 x 10^6^ cells/ml. The cell viability should be at least 70% for optimal processing. These cell suspensions were loaded into the 10X Genomics Chromium instrument, where approximately 10,000 cells from each sample were captured. The library was then processed using the 5’ whole transcriptome gene expression workflow and paired full-length TCR sequencing^65^.

scRNA-seq analyses followed our general procedure described in the “**10X scRNA-seq/scTCR-seq data analyses**” section below. Cell typing was performed by referring to cell markers from Hurskainen *et al*^66^. The typing for tumor *vs.* normal epithelial cells was confirmed by checking whether the corresponding sequencing reads contained *Kras*^G12D^ and *Trp53*^R172H^ mutations (**Sup. File 3**). MHC-binding peptides were predicted by NetMHCpan v4 and NetMHCIIpan v4, for the MHCs of the 129Sv mice (H-2-Kb, H-2-Db, and H-2-IAb).

### ICI irAE patient cohort and generation of genomics data

The study was performed following protocols approved by the Institutional Review Board of the University of Texas Southwestern Medical Center(“Biospecimen procurement for immunologic correlates in cancer”, IRB number is 082015-053). Patients provided written, informed consent for additional data collection, presentation, and publication. Clinical, radiographic, and laboratory data were collected from electronic health records (Epic Systems, Verona, Wisconsin). Due to the complexity and difficulty of diagnosing and characterizing irAE, each case was reviewed independently by two clinicians experienced in ICI administration and monitoring, with any differences resolved through third-party adjudication. Detailed patient characteristics are described in **Sup. Table 6**.

As previously described^67^, peripheral blood samples were examined from the patients at pre-ICI baseline, 2 weeks, 6 weeks, 3 months, every 3 months thereafter, and after initiation of steroids for irAE treatment, though sample collection or data generation failed for some patients at some time points. We also examined serial samples (baseline, 2 weeks) from two healthy controls. Samples were centrifuged at 3000 rpm at 4°C for 15 min to obtain plasma. Peripheral blood mononuclear cells (PBMCs) were isolated from samples using density gradient centrifugation in Ficoll-Paque Plus Media (Fisher Scientific, Waltham, Massachusetts).

For TCR-seq, DNA quality was assessed using the Agilent TapeStation instrument with the Genomic DNA Screentape reagent kit by Agilent Technologies (Catalog #5067-5365 and #5067-5366). DNA concentration was determined using the Quant-iT™ PicoGreen dsDNA Assay kit by Invitrogen (Catalog #P7589) and a PerkinElmer plate reader (PerkinElmer Victor X3, 2030 Multilabel Reader). The "immunoSEQ hsTCRB v4b" Kit (Product Code: ISK10050) by Adaptive Biotechnologies and the "Qiagen Multiplex PCR Plus" Kit (Product Code: 206152) were used for TCR-seq library generation. The procedure followed the manufacturers’ instructions. Kit information is available at www.adaptivebiotech.com/adaptive-immunosequencing. The workflow included PCR plate layout design, two PCR amplifications, library pooling and purification, and library quality and quantity assessment. Specific primers, incorporating a proprietary mix of V- and J-genes, were used in the first PCR amplification to amplify all potential TCRβ rearrangements, along with reference gene primers. The second PCR amplification added unique DNA barcodes and Illumina adapters to each replicate. Following this, libraries were pooled and purified using the provided kit beads. Finally, library quantity was measured using the PicoGreen method, and quality was verified on an Agilent 2100 Bioanalyzer instrument with the Agilent DNA 1000 kit (Agilent Technologies, Catalog# 5067-1504). Additionally, adapter ligation efficiency was confirmed through qPCR quantification using the Kapa Library Quant Kit by Roche (Catalog # 7960336001). All samples were sequenced on the Illumina NextSeq 550 sequencer platform with 150 cycles sequencing kits, following the Adaptive Biotechnologies sequencing protocol. Raw sequencing data were transferred to Adaptive Biotechnologies, where the data analysis was carried out by their dedicated data analysis team. As Adaptive could not provide matched α-β TCR chain sequences, pMTnet-omni was applied to β chain TCR data only to predict TCR-pMHC binding, as was done in **Fig. 1f**.

Monitoring of cytokine and chemokine levels was performed using Bio-Plex Pro Human Chemokine 40-plex Panel (Bio-Rad Laboratories, Hercules, California) using a Luminex 200 System. Bio-Plex Manager 6.1 software was used for data analysis. Concentrations of cytokines and chemokines (pg/mL) were determined on the basis of the fit of a standard curve for mean fluorescence intensity versus pg/mL. These cytokines are stable over time in healthy controls not receiving ICI^68^.

### Calculation of the irAE risk score

We devised an irAE risk score by integrating the pMTnet-omni binding prediction results with the TCR clonal expansion information. The risk score is computed for each irAE and as a normalized percentage of TCRs that target the putative auto-antigens from each organ affected by the irAE of interest, among the TCR clones with the highest clonal expansions. It can be intuitively understood as a measurement of “correlation” between TCR clonal expansions and binding predictions akin to a Fisher’s exact test, where we dichotomize TCRs by their clonal sizes and auto-antigenic pMHCs by the pMTnet-omni binding predictions. Technically, given a sequence of TCR clonotypes in decreasing order of clonal sizes: *TCR_1_, …, TCR_T_*, and a set of pMHCs: *pMHC_1_, …, pMHC_p_*, from the auto-antigens of the organ affected by the irAE and presented by the patient-specific HLA alleles, we first compute, *via* pMTnet-omni, their rank%s *r_11_,…, r_1P_,…,r_t1_,…,r_tP_,…,r_T1_,…,r_TP_*. We then select a cutoff *C* such that the percentage of binding TCR-pMHC pairs is calculated only based on the top *C* most expanded TCRs. The risk score is then computed as

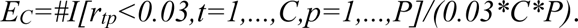

In our analysis, the final risk score is found *via*

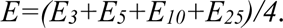

### Pre-processing of the COVID-19 vaccination dataset

The scRNA-seq/TCR-seq and the patients’ metadata were obtained from Curtis *et al*^46^. scRNA-seq analyses followed our general procedure described in the “**10X scRNA-seq/scTCR-seq data analyses**” section below. SARS-CoV-2 viral protein sequences were collected from UniProt (www.uniprot.org), including Replicase polyprotein 1ab (P0DTD1), Spike glycoprotein (P0DTC2), Envelope small membrane protein (P0DTC4), Membrane protein (P0DTC5), Nucleoprotein (P0DTC9), *ORF7a* (P0DTC7), *ORF8* (P0DTC8), *ORF9b* (P0DTD2), *ORF3a* (P0DTC3), *ORF6* (P0DTC6), *ORF7b* (P0DTD8), *ORF9c* (P0DTD3), *ORF3b* (P0DTF1), *ORF10* (A0A663DJA2), *ORF3c* (P0DTG1), and *ORF3d* (P0DTG0). Protein peptides presented by class I and class II MHCs were predicted by netMHCpan and netMHCIIpan. HLA allele information of each patient was obtained from the original authors of this dataset.

### Pre-processing of the 10X scRNA-seq/scTCR-seq/antigen-affinity data

scRNA-seq analyses followed our general procedure described in the “**10X scRNA-seq/scTCR-seq data analyses**” section below. TCRs having TRUE attributes for “is_cell”, “high_confidence”, “full_length”, and “productive”, and having only 1 CDR3α sequence and 1 CDR3β sequence were retained for the following analysis. Antigen affinity is measured by 10X’s feature-barcoding technology and the output is a UMI count. For each TCR clonotype and each pMHC, the UMI counts of T cells in this clonotype were averaged as the binding score for that pMHC. Then the pMHC receiving the maximum averaged UMI counts, which must also be larger than 5, was assigned as the binding pMHC for this TCR clonotype.

### Pre-processing of the Visium-TCR-seq data

We used arcasHLA^48^ to perform HLA typing on the transcriptomic component of the Visium-TCR data. The identification of the TAAs and pMHCs has been described in the result section. We processed the TCR-sequencing component of the Visium-TCR-seq data with MiXCR^69^ to obtain the β chain TCR sequences. As Visium-TCR-seq is not performed at the single cell resolution, we cannot obtain matched α-β TCR chain sequences. Therefore, pMTnet-omni was applied to β chain TCR data only to predict their binding towards TAA pMHCs, as was in **Fig. 1f**.

### Correlation between TAA expression and clonal expansion of TAA-specific TCRs

Based on our results on classifying spots into tumor, stromal/immune, and blood spots, we identified the tumor-stroma/immune interface by examining the surrounding 4 layers of spots of each sequencing spot. If the type of at least one of the surrounding spots is different from the type of the spot under examination, this position is classified as a part of the tumor-stroma/immune interface (**Fig. 5d**). To compute the correlation between the gene expression levels of a TAA at a sequencing spot and the clonal expansion of the nearby TCRs targeting that TAA, we first leveraged the prediction results of pMTnet-omni and retained TCR clones that were predicted as binding with the rank% cutoff being 3% in each sequencing spot. Then the clonal expansion of nearby TCRs for a particular spot was calculated as the sum of the clonal sizes of all binding TCRs in its surrounding 3 layers of spots. After the adjustment was done, the correlation was found using Spearman’s rank correlation coefficient.

### Pre-processing of PDB structures of TCR-pMHC complexes

Sequence information on the α chain, β chain, MHC, and the presented peptide, as well as the PDB file for each TCR-pMHC complex, was downloaded from tcr3d.ibbr.umd.edu/downloads. IgBlast (www.ncbi.nlm.nih.gov/igblast) was used to annotate the TCR chains. Then we computed the distances between each amino acid residue of CDR3ɑ/CDR3β to all amino acid residues of the pMHC and took the minimum of all these distances for each CDR3 residue. Alanine scanning as well as “all other AA” scanning were implemented by simply replacing a given amino acid residue on CDR3ɑ/CDR3β with each one of the other 19 amino acids.

### Execution of benchmark software

All calculations based on the four variations of ERGO-II were carried out using the authors’ web tool: tcr2.cs.biu.ac.il/home. ATM-TCR (github.com/Lee-CBG/ATM-TCR) and ImRex (github.com/pmoris/ImRex) were installed according to their respective documentation. When comparing ERGO-II with pMTnet-omni, we only retained human pMHC-TCR pairs due to the limit of ERGO-II to human pairs only. As ATM-TCR is only applicable for peptides presented by class I MHCs, we only considered class I pMHC-TCR pairs in their comparison against pMTnet-omni. Similarly, we only kept human pairs for the benchmark against ImRex due to its limitation. Besides, ATM-TCR and ImRex only accept CDR3β as input. For all benchmark software, default parameters were used during the benchmark study.

### 10X scRNA-seq/scTCR-seq data analyses

All scRNA-seq/scTCR-seq data involved in this study were processed by the standard Cellranger-multi pipeline. The scRNA-seq data was analyzed with Seurat (v4.0.6). Cells with detected genes larger than 200, mitochondrial counts fewer than 10%, and total molecules within a cell larger than 500 were retained. Genes detected in more than 10 cells were retained. Then the count data was normalized by *NormalizeData* function with default parameters. Highly variable features were identified by *FindVariableFeatures* with default parameters. After that, datasets from all samples were integrated by *SelectIntegrationFeatures*, *FindIntegrationAnchors*, and *IntegrateData*. Principal component analysis was run with the first 100 principal components with *RunPCA*. Neighbors and clusters of cells were identified by *FindNeighbors (first 100 dimensions)* and *FindClusters (resolution=1)*. Uniform manifold approximation and projection (UMAP) was run with *RunUMAP (the first 100 dimensions)*.

### Statistical analyses

Computations were mainly performed in the R (3.6.3 and 4.1.3, 4.1.1) and Python (3.7 and 3.8) programming languages. All statistical tests were two-sided unless otherwise described. For all boxplots appearing in this article, box boundaries represent interquartile ranges, whiskers extend to the most extreme data point, which is no more than 1.5 times the interquartile range, and the line in the middle of the box represents the median. UMAP was created by Seurat v4.0.6. The heatmap was created by ComplexHeatmap v2.10.0. Gene ontology analysis was performed by clusterProfiler v4.2.2. The Circos plots were created by the R library circlize v0.4.15. The mixed effect models were fitted using the R library lme4 v1.1.32 with TCR-irAE risk scores being the fixed effect and random intercepts and slopes within each irAE. The fast gene set enrichment analysis was performed using the R package fgsea v1.24.0. The 3D waterfall plot was created by the R packages ggplot2 v3.4.4, rgl v1.2.1, and devoutrgl v0.1.0. PyMOL 2.5.7 was used to render the visualization of the 3D structure of the TCR-pMHC complex.

### Data availability

For reviewers, please use this login to access our archived data on DBAI: username, password is: reviewer123. We limited access to these data during the review phase due to confidentiality reasons, and will openly release when the paper is accepted.

**Sup. Table 1** documents the TCR sequence datasets that were used for training the V and CDR3 VAEs. The p-MHC binding data that we used for training and validating the pMHC encoder are available at www.iedb.org/database_export_v3.php, and services.healthtech.dtu.dk/suppl/immunology/NAR_NetMHCpan_NetMHCIIpan/. **Sup. Table 2** documents the TCR-pMHC pairing datasets that we collected for training and validation of the final stage of pMTnet-omni. The mouse tumor progression scRNA/TCR-seq data can be accessed on the Database for Actionable Immunology (DBAI) website (dbai.biohpc.swmed.edu)^9,10,70^, in the scRNA data table with accession numbers from scRNA_00000000000001 to scRNA_00000000000014. For the irAE cohort, patient clinical information can be found in **Sup. Table 6**, TCR-sequencing data can be accessed in the receptor data table of DBAI with accession numbers from Receptor_00000000000001 to Receptor_00000000000113, and the cytokine profiling data can be found in FigShare under the DOI: https://doi.org/10.6084/m9.figshare.24570136.v2. The COVID-19 vaccination data (processed scRNA-seq, TCR-seq, and metadata) were obtained from the original publication^46^. The Visium-TCR-seq data were obtained from www.ncbi.nlm.nih.gov/sra/?term=PRJNA742564 and www.ncbi.nlm.nih.gov/geo/query/acc.cgi?acc=GSE179572. The four 10X datasets (used in **Fig. 6**) with multiplexed scRNA-seq, scTCR-seq, and antigen affinity information were downloaded from the 10X Genomics website (more information: www.10xgenomics.com/resources/document-library/a14cde). The PDB structure data were downloaded from tcr3d.ibbr.umd.edu/downloads.

### Code availability

For reviewers, also for confidentiality reasons here, please check the attached compressed file (software.tar.gz) containing the source codes of pMTnet-omni and associated files needed for executing pMTnet-omni.

The pMTnet-omni tool is provided as a web service on the Database for Actionable Immunology (DBAI) website^9,10,70^, under the “Online Tools” tab: dbai.biohpc.swmed.edu/pmtnet. Tutorials are provided for users to understand input/output format and model parameters.

## ACKNOWLEDGEMENTS

We acknowledge Dr. Jason Y. Park for his valuable suggestions on the irAE section of our work. This study was supported by the National Institutes of Health (NIH) [5R01CA258584/TW, XW, R01CA234629-01/JZ, 1U01AI156189/DG, R38HL150214/MG], Cancer Prevention Research Institute of Texas [RP190208/TW, RP230363/TW, JH, AR], Khalifa Scholar Award [JZ], MD Anderson Lung Cancer Moon Shot Program [JZ], American Cancer Society-Melanoma Research Alliance Team Award [MRAT-18-114-01-LIB/DB], and V Foundation Robin Roberts Cancer Survivorship Award [DT2019-007/DB].

## AUTHOR CONTRIBUTIONS

Y.H., Y.Y, X.K., Y.H., O.X. contributed to all bioinformatics analyses. Y.T, D.G., C.C., J.Z. generated the mouse tumor progression datasets. F.J.F., M.S.I., Y.X., C.L., I.R., C.Z., M.E.G., J.E.D., J.H., S.R., D.H., Y.G., D.E.G. collected the irAE patient cohort and performed all experimental assays. D.Y., J.L., S.Y. curated patient information, especially the clinical variables, for the irAE cohort. M.Z., K.P., C.Y., A.R. performed validation experiments for TCR-pMHC pairs predicted by pMTnet-omni. F.W. created the DBAI/pMTnet-omni web service. X.W., J.H. provided critical input on the manuscript. T.W. supervised the whole study. All authors wrote the manuscript.

