## Supplementary figures and images for "pan-MHC and cross-Species Prediction of T Cell Receptor-Antigen Binding"

### Sup. Fig. 1

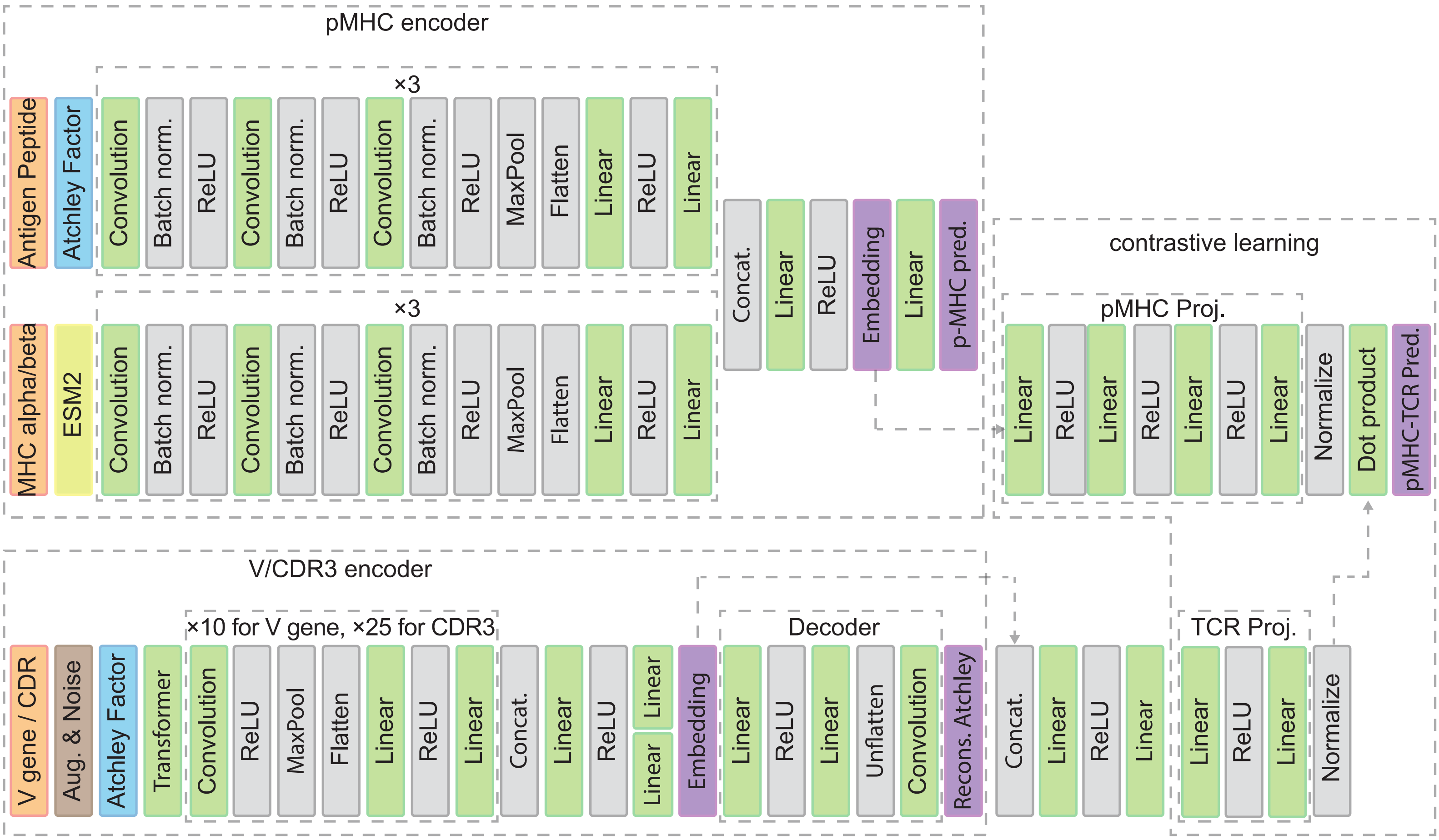

### Sup. Fig. 2

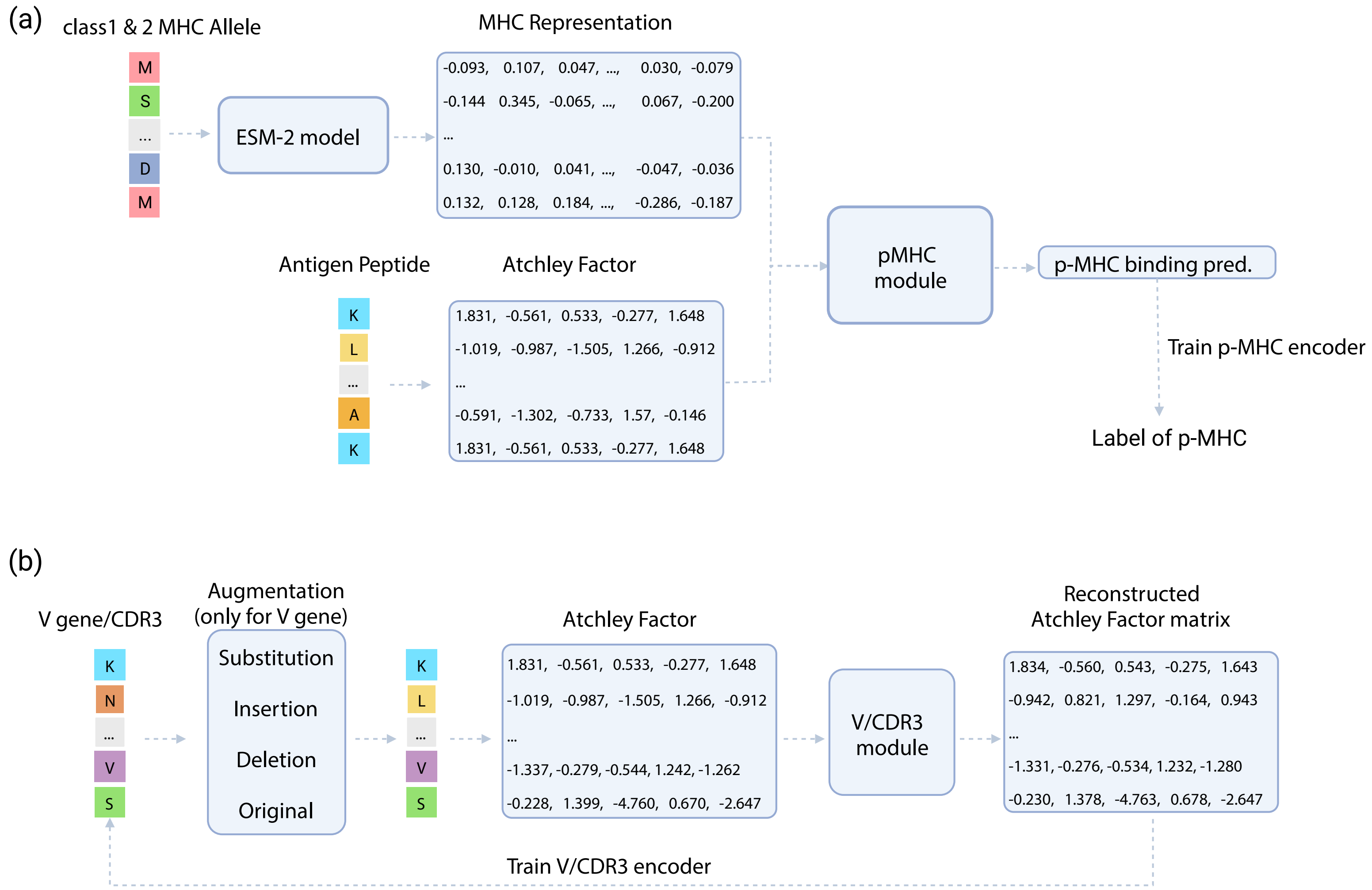

### Sup. Fig. 3

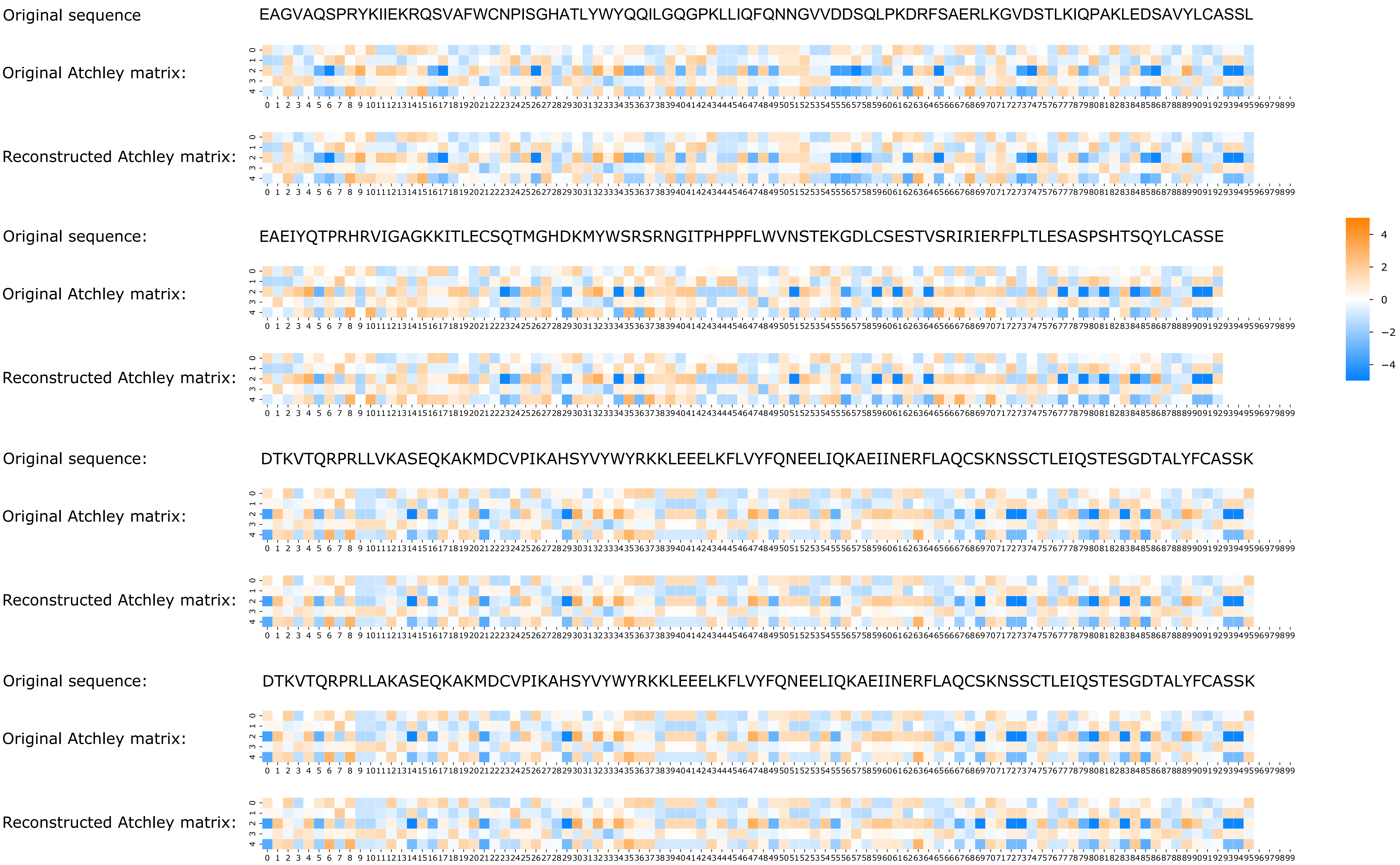

### Sup. Fig. 4

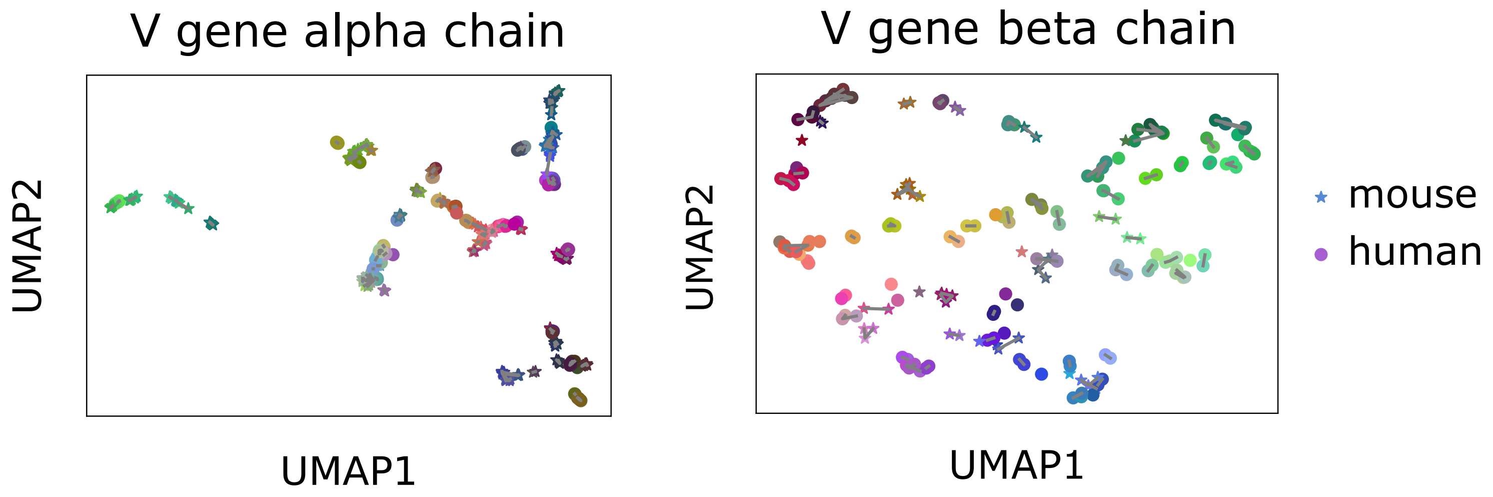

### Sup. Fig. 5

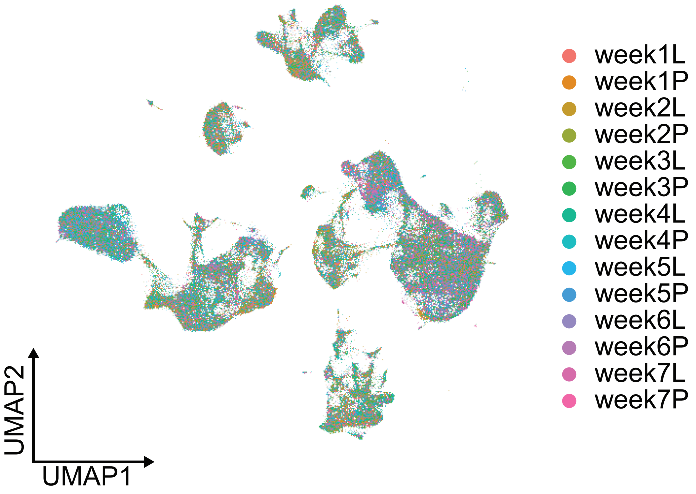

### Sup. Fig. 6

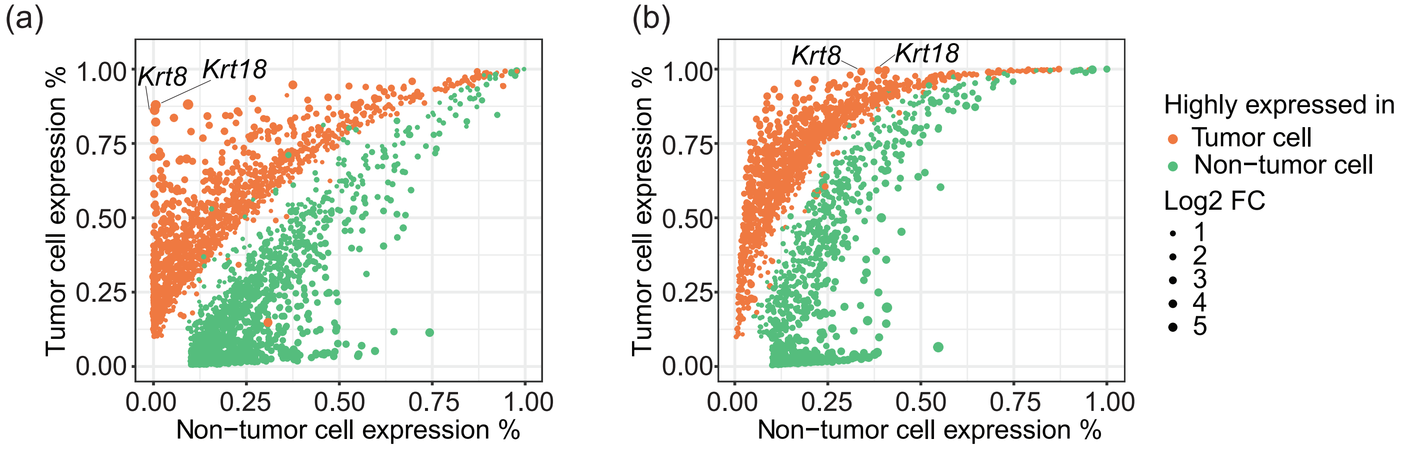

### Sup. Fig. 7

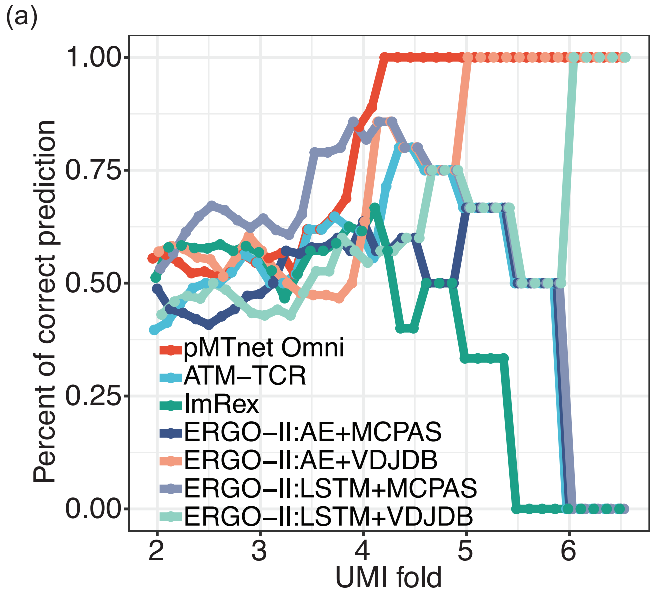

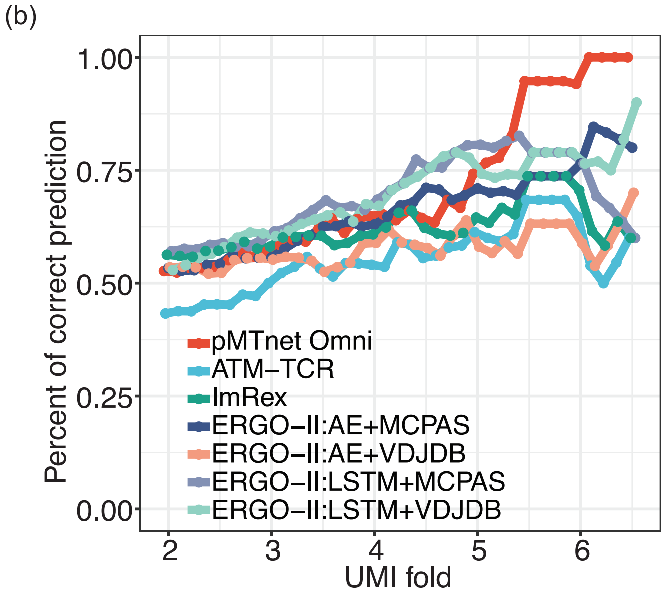
