## Supplementary material for "pan-MHC and cross-Species Prediction of T Cell Receptor-Antigen Binding": Sup. File 1

**Supplementary File 1: Additional details of model architecture and training procedure**

**Peptide Atchley Factor representation**

The peptide amino acids (AAs) were first converted into numerical vectors according to the Atchley Factor[^1^](https://sciwheel.com/work/citation?ids=5389716&pre=&suf=&sa=0). Peptides shorter than 30 AAs were padded to 30 AAs and the padded amino acids were converted to 0 in the numerical vectors. Peptides longer than 30 AAs were truncated to 30 AAs. Then the numerical vectors of peptides were fed into the pMHC-omni model.

**MHC representation**

Human MHC allele sequences were collected from the IPD-IMGT/HLA Database (www.ebi.ac.uk/ipd/imgt/hla). Mouse MHC allele sequences were collected from the UniProt Database (www.uniprot.org) and NCBI (www.ncbi.nlm.nih.gov). The ESM2 650M model[^2^](https://sciwheel.com/work/citation?ids=14536375&pre=&suf=&sa=0) was employed to obtain the representation of MHC alleles. The MHC protein sequences were padded to 380 AAs. Then the MHC sequences were fed into the ESM2 model to obtain MHC protein embeddings.

**peptide-MHC binding prediction**

To train the p-MHC model, the 30-dim final concatenated p-MHC embeddings were linearly transformed into a 3-dim predictor layer. Mean squared error (MSE) loss and cross-entropy loss were used for binding affinity training data (BA, continuous output) and eluted ligands training data (EL, binary output), respectively. Two dims of the predictor layer were input into the cross-entropy loss and another one dim of the predictor layer was input into the MSE loss.

**V gene and CDR3 encoders**

The V gene and CDR3 AA sequences were also converted to numerical vectors according to the Atchley Factor. V gene alleles and amino acid sequences were downloaded from IMGT. 21 human Vα alleles (TRAV11-1*01, TRAV14-1*01, TRAV14-1*02, TRAV15*01, TRAV28*01, TRAV28*02, TRAV29/DV5*03, TRAV3*02, TRAV32*01, TRAV33*01, TRAV33*02, TRAV35*03, TRAV37*01, TRAV46*01, TRAV8-5*01, TRAV8-6-1*01, TRAVA*01, TRAVA*02,TRAVB*01, TRAVB*02, TRAVC*01), 26 human Vβ alleles (TRBV10-1*03, TRBV12-1*01, TRBV12-2*01, TRBV16*02, TRBV22-1*01, TRBV22/OR9-2*01, TRBV24/OR9-2*02, TRBV24/OR9-2*03, TRBV25/OR9-2*02, TRBV26*01, TRBV26/OR9-2*01, TRBV26/OR9-2*02, TRBV3-2*01, TRBV3-2*02, TRBV3-2*03, TRBV30*03, TRBV5-2*01, TRBV8-1*01, TRBV8-1*02, TRBV8-2*01, TRBV8-2*02, TRBVA*01, TRBVA*02, TRBVA/OR9-2*01, TRBVB*01, TRBVB*02), 35 mouse Vα alleles (TRAV11N*01, TRAV12-4*01, TRAV12-4*02, TRAV12-4*03, TRAV13-3*03, TRAV15-3*01, TRAV15-3*02, TRAV15D-3*01, TRAV15D-3*02, TRAV15N-3*01, TRAV19*02, TRAV20*01, TRAV20*02, TRAV22*01, TRAV22*02, TRAV23*02, TRAV3-2*01, TRAV3-2*02, TRAV3D-2*01, TRAV3D-2*02, TRAV3N-2*01, TRAV4-1*01, TRAV4-1*02, TRAV4D-2*01, TRAV4D-2*02, TRAV5-2*01, TRAV5-2*02, TRAV5-3*01, TRAV5D-2*01, TRAV5D-2*02, TRAV5D-3*02, TRAV5N-2*01, TRAV5N-3*01, TRAV6-7/DV9*05, TRAV9N-1*01), and 12 mouse Vβ alleles (TRBV11*01, TRBV12-3*01, TRBV18*01, TRBV22*01, TRBV24*02, TRBV25*01, TRBV27*01, TRBV28*01, TRBV6*01, TRBV7*01, TRBV8*01, TRBV9*01) contained stop codons (asterisk) in the middle of their sequences and were dropped. Due to the small numbers of V genes/alleles available for deep learning, the V gene sequences were augmented before being converted to numerical vectors. For each V gene sequence, there was a 25% probability of adding a random AA to the original sequence, a 25% probability of switching the AA at any position to a random one of the other 19 possible AAs, a 25% probability of deleting an AA at any position, and a 25% probability of remaining unchanged. The augmentation process was repeated 10,000 times. The V gene AA sequence was padded or cut to at most 100 AAs. The CDR3 AA sequence was padded or cut to at most 25 AAs. The numerical vectors of the padded AAs were masked and then fed into the VAE encoders. The loss function of the V gene VAE models and CDR3 VAE models was the sum of the reconstruction loss and Kullback-Leibler(KL) divergence loss with more emphasis on the reconstruction loss. The weights of KL divergence loss in Vα, Vβ, CDR3α, and CDR3β models were 1E-5, 1E-4, 1E-5, and 1E-5.

**pMTnet-omni contrastive learning**

In the final pMHC-TCR contrastive learning stage, the embeddings of pMHC were fed into the projection layer, where the linear layers’ out-feature sizes were 40, 50, 60, and 70. The TCR (Vα, Vβ, CDR3α, and CDR3β) embeddings were first concatenated and the linear layer’s out-feature sizes were all 70. The temperature parameter in the contrastive learning was set to 0.1. Supervised contrastive loss[^3^](https://sciwheel.com/work/citation?ids=14754084&pre=&suf=&sa=0) was employed, and the training process was repeated 3 times with 3 random seeds: 50, 51, and 52. The model parameters were saved at the 65^th^ epoch.

**Optimization algorithm**

The pMHC, Vα, Vβ, CDR3α, CDR3β, and final pMHC-TCR models were trained separately. The Adam algorithm was used as the optimizer for training the pMHC, V gene, and CDR3 models with learning rates set to 1E-3, 1E-4, and 5E-4. To reduce memory usage and accelerate computation, automatic mixed precision in PyTorch was used in the training of pMHC, V gene, and CDR3 models. The AdamW algorithm was used as the optimizer for training the pMHC-TCR model with a learning rate set to 5E-4 and a weight decay set to 1E-4. A linear scheduler with warmup was used to reduce learning rate for the pMHC-TCR model.

[Bibliography](https://sciwheel.com/work/bibliography)

[1.	Atchley, W.R., Zhao, J., Fernandes, A.D., and Drüke, T. (2005). Solving the protein sequence metric problem. Proc Natl Acad Sci USA *102*, 6395–6400. 10.1073/pnas.0408677102.](https://sciwheel.com/work/bibliography/5389716)

[2.	Lin, Z., Akin, H., Rao, R., Hie, B., Zhu, Z., Lu, W., Smetanin, N., Verkuil, R., Kabeli, O., Shmueli, Y., et al. (2023). Evolutionary-scale prediction of atomic-level protein structure with a language model. Science *379*, 1123–1130. 10.1126/science.ade2574.](https://sciwheel.com/work/bibliography/14536375)

[3.	Khosla, P., Teterwak, P., Wang, C., Sarna, A., Tian, Y., Isola, P., Maschinot, A., Liu, C., and Krishnan, D. (2020). Supervised Contrastive Learning. arXiv. 10.48550/arxiv.2004.11362.](https://sciwheel.com/work/bibliography/14754084)
