## Supplementary material for "pan-MHC and cross-Species Prediction of T Cell Receptor-Antigen Binding": Sup. File 2

**Sup. File 2: Curation of TCR-antigen pairing data**

**Dataset selection**

We searched for scientific literature that was published before our data lock date of Dec, 1^st^, 2022, which contains data on human or mouse TCR-T cell antigen pairing data. We curated the pairing data, if available, from these scientific publications. We consider both CD8^+^ T cell/MHC class I pairings and CD4^+^ T cell/MHC class II pairings. The pairing information is derived from a variety of technologies, such as tetramer assays, high throughput sequencing-based screening approaches, *etc*. Several of our data sources are pre-existing TCR-antigen pairing databases, such as IEDB. We included a study only when this study contained pairing data of matched α chains and β chains for at least some of the pairing records.

**Nomenclature harmonization**

We harmonized the naming conventions of all the pairing datasets (N=70) we collected from the previous step. Specifically, the datasets were examined individually so that only 8 columns were retained: TRA_V (the name of the V$\alpha$ gene of a T cell receptor), CDR3_ALPHA (the Complementarity Determining Region 3 of the $\alpha$ chain), TRB_V (the name of the V$\beta$ gene of a T cell receptor), CDR3_BETA (the Complementarity Determining Region 3 of the $\beta$ chain), EPITOPE (the peptide presented by major histocompatibility complex), MHC (the major histocompatibility complex), pMHC_SPECIES (human or mouse), and TCR_SPECIES (human or mouse). We also made sure that the values of TRA_V and TRB_V start with “TRAV” and “TRBV”, respectively, that a human MHC starts with “HLA-”, and that a mouse MHC starts with “H-2”. V allele names were corrected according to IMGT (www.imgt.org). Human MHC allele names were corrected according to the IPD-IMGT/HLA Database (www.ebi.ac.uk/ipd/imgt/hla). "DPA1*01:01", "A*24:01", "B*07:01", "B*44:01" were replaced with "DPA1*01:03", "A*24:02", "B*07:02", "B*44:02" because their sequences contained errors and they were deleted in the IPD-IMGT/HLA database. "B*12" was replaced with "B*44:02" according to prior reports[^1,2^](https://sciwheel.com/work/citation?ids=15651928,15651929&pre=&pre=&suf=&suf=&sa=0,0). The values for EPITOPE, MHC, pMHC_SPECIES, and TCR_SPECIES are present for all data records. We allow missingness in the TRA_V, CDR3_ALPHA, TRB_V, and CDR3_BETA attributes. But we only keep a record if at least one of the CDR3_ALPHA and CDR3_BETA attributes is present.

**Quality control**

This step of data cleansing validates all gene IDs and amino acid symbols. To start with, some obvious typos such as the usage of an exotic symbol like “–” in place of a “-” were corrected. Leading and trailing whitespaces were stripped, while missing colons and asterisks were added. Then, three resources were loaded for the purpose of data verification: the one-letter codes for the 20 amino acids; the mouse MHC gene and allele names found on UniProt and IEDB; and the standardized names of the V genes on the $\alpha$ chains and the $\beta$ chains of human and mouse T cell receptors recorded from UniProt. We first checked if each of the amino acids in CDR3_ALPHA, CDR3_BETA, and EPITOPE columns was one of the 20 amino acids. If there was any symbol outside of the 20 letters in an EPITOPE, we filtered out this record. If there was any symbol outside of the 20 letters in CDR3_ALPHA and CDR3_BETA, we set the records to blank (as if CDR3_ALPHA or CDR3_BETA is missing). CDR3_ALPHA and CDR3_BETA having at least 5 amino acids were retained. If we cannot find the MHC gene name for any record, we will remove this record. If we cannot find the V gene names for any record, we set the V gene names to blank (as if missing). Pseudo V gene alleles are also treated as missing values.

For human and mouse MHCs, we further created two MHC_ALPHA and MHC_BETA attributes. For Class I human MHCs, the MHC allele names were copied to their corresponding MHC_ALPHA column. The MHC_BETA fields were filled with “human_microglobulin” which was retrieved from the UniProt database(www.uniprot.org). For Class II human HLAs, we copied the $\alpha$ allele and the $\beta$ allele to MHC_ALPHA and MHC_BETA, respectively. But for the HLA-DRs, if only the DRB allele names exist, we copied them to the MHC_BETA column and fill the MHC_ALPHA records with “DRA*01:01”. There are only two different coding alleles for HLA-DRA, with minimum differences. HLA-DRA is usually not typed in most HLA typing assays, and thus not available to us for most cases. Similarly, the values of Class I mouse MHCs were copied to the corresponding MHC_ALPHA column. The MHC_BETA fields were filled with “mouse_microglobulin”. The $\alpha$ allele and the $\beta$ allele of mouse class II MHC heterodimers have fixed combinations of the $\alpha$ allele and the $\beta$ allele (unlike human class II alleles where $\alpha$ alleles can be paired with different $\beta$ alleles). So the class II mouse MHCs were simply copied to both MHC_ALPHA and MHC_BETA columns with an additional “_alpha” or “_beta” attached at the end to differentiate. The sequences of the $\alpha$ alleles and the $\beta$ alleles of the mouse class II MHC heterodimers were extracted from the UniProt and NCBI databases.

For validation of V gene names, we found that there were two nomenclature systems of V genes used in our datasets: the IMGT nomenclature and the naming system adopted by Arden *et al*[*^3^*](https://sciwheel.com/work/citation?ids=6159421&pre=&suf=&sa=0). The tables containing correspondence between nomenclatures provided by IMGT were used to map all V gene names to IMGT nomenclature. Another subtle nuisance was that if a record of a V gene did not possess the allele level designation, we appended “*01” at the end of this gene name, which is usually the reference allele of this V gene.

**Training-validation split**

The TCR-antigen pairing records from the steps above are checked again and we only keep records (1) with complete EPITOPE, MHC_ALPHA, MHC_BETA, pMHC_SPECIES, and TCR_SPECIES information, and (2) if at least one of CDR3_ALPHA and CDR3_BETA is present. Our collected datasets were split into training and validation according to their data quality and sample sizes. In general, we prioritize high-quality datasets for validation purpose and larger datasets (such as those from high-throughput screening technologies or database dumps) for training purpose. After data cleaning, a total of 19 datasets were split into the training subset and 50 datasets were split into the validation subset (**Sup. Table 2**).

In the validation subset, only TCR-pMHC binding pairs (positive pairs) with complete information (V gene alleles and CDR3 of both $\alpha$ and $\beta$ chains) were retained. Negative binding pairs were manually created by substituting the TCRs in these positive binding pairs with randomly sampled background TCRs (**Sup. Table 1**). Among these constructed negative validation pairs, some coincided with positive binding pairs and they were dropped. The negative validation pairs were synthesized to be 5 times as many as the positive validation pairs. Finally, there were 812 positive and 4,060 negative pairs in the validation cohort.

We then processed the training data. Both complete and incomplete binding pairs were retained as positive training pairs in order to achieve a larger sample size. Overlapping pairs that also appeared in the validation cohort were dropped. Here, “different pairs” were defined as two pairs, where, for complete data slots (non-missing in both pairs), at least one slot among TRA_V, TRB_V, CDR3_ALPHA, CDR3_BETA, TCR_Species, EPITOPE, MHC_ALPHA, MHC_BETA, MHC_species, is different across the two pairs. Finally, there were 110,390 unique positive pairs. Due to the small sample sizes of the human CD4^+^ T cell/MHC class II pairs and mouse TCR-MHC pairs, positive human CD4^+^ T cell/MHC class II pairs and positive mouse TCR-MHC pairs were upsampled 10 times. Negative training pairs were manually created by substituting the TCRs in the positive training pairs with randomly sampled background TCRs, as above. For the set of negative pairs created for each positive pair, the pattern of missingness was kept the same. For example, if the positive pair is missing TRA_V, all the negative pairs created from this positive pair will miss TRA_V. Among these constructed negative training pairs, some coincided with the positive binding pairs in the training/validation subset or the negative pairs in the validation subset, and they were again dropped. In each Epoch, the negative training pairs were synthesized to be 50 times as many as the positive pairs in the training subset.

[**Bibliography**](https://sciwheel.com/work/bibliography)

[1.	Weekes, M.P., Wills, M.R., Mynard, K., Carmichael, A.J., and Sissons, J.G. (1999). The memory cytotoxic T-lymphocyte (CTL) response to human cytomegalovirus infection contains individual peptide-specific CTL clones that have undergone extensive expansion in vivo. J. Virol. *73*, 2099–2108. 10.1128/JVI.73.3.2099-2108.1999.](https://sciwheel.com/work/bibliography/15651928)

[2.	Schiffer, C.A., O’Connell, B., and Lee, E.J. (1989). Platelet transfusion therapy for alloimmunized patients: selective mismatching for HLA B12, an antigen with variable expression on platelets. Blood *74*, 1172–1176.](https://sciwheel.com/work/bibliography/15651929)

[3.	Arden, B., Clark, S.P., Kabelitz, D., and Mak, T.W. (1995). Human T-cell receptor variable gene segment families. Immunogenetics *42*, 455–500.](https://sciwheel.com/work/bibliography/6159421)
