## Supplementary material for "pan-MHC and cross-Species Prediction of T Cell Receptor-Antigen Binding": Sup. File 3

**Supplementary File 3: Additional analyses and results**

**Cell typing of scRNA-seq data from the 344SQ/129Sv mouse model**

The scRNA-seq data of the primary and metastasis tissue samples were integrated and processed with a standard pipeline using Seurat. Eleven types of cells, including epithelial cells, were annotated based on established cell markers[^1–3^](https://sciwheel.com/work/citation?ids=11169341,9507947,10054525&pre=&pre=&pre=&suf=&suf=&suf=&sa=0,0,0) (**Sup. File 3 Fig. 1a**). To differentiate tumor epithelial cells from normal epithelial cells, scRNA-seq reads with *Kras*^G12D^/*Tp53^R172H^* mutations and reads with wild type *Kras*/*Tp53* nucleotides in the same position were counted. Based on the distribution of these single cells carrying mutant *vs* wild-type reads in the UMAP plot (**Sup. File 3 Fig. 1b-e**), we designated which clusters of epithelial cells are tumor *vs* normal epithelial cells.

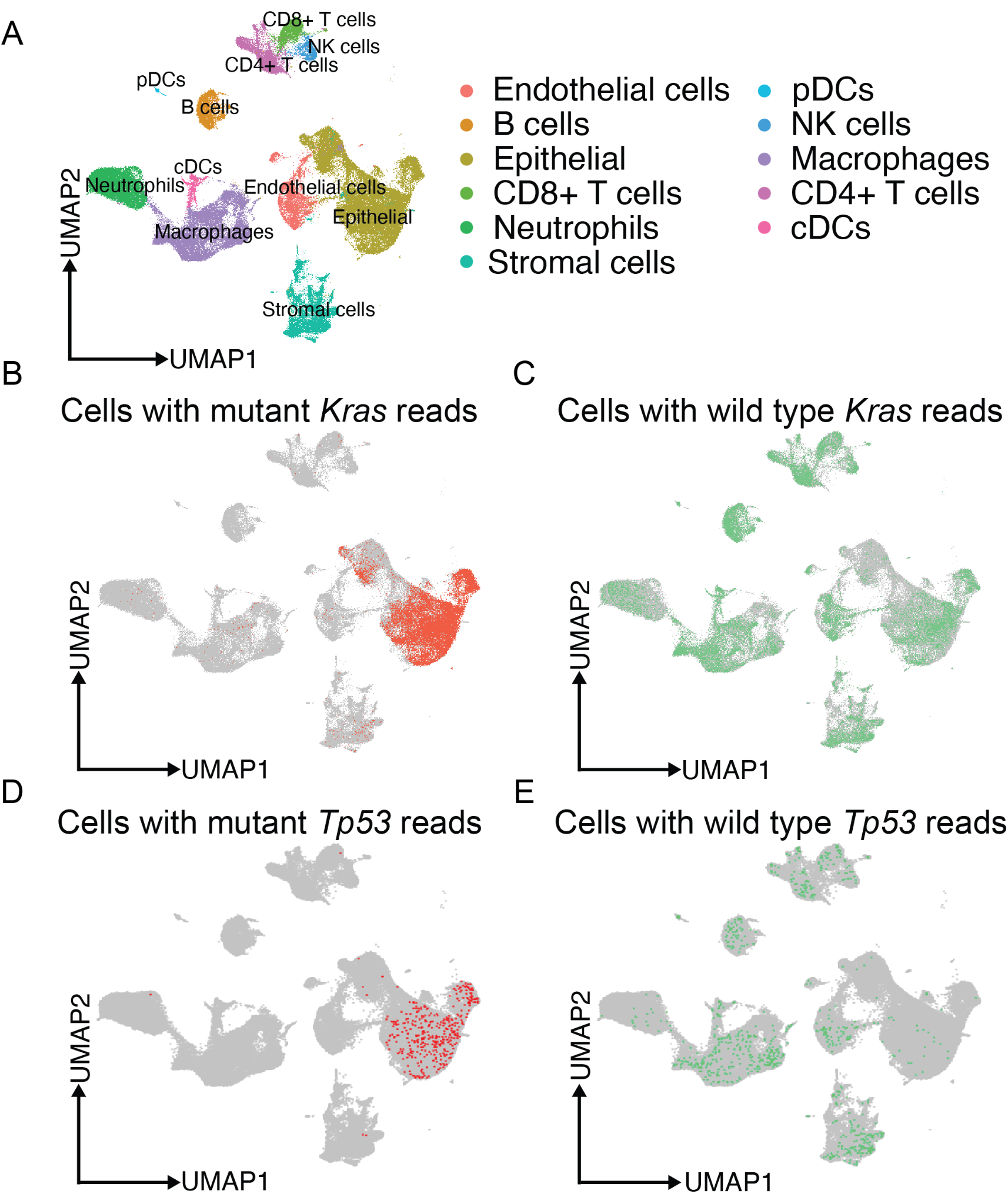

***Sup. File 3 Fig. 1*** *Cell typing for the 344SQ/129Sv mouse model. (a) UMAP showing the annotated cell types based on gene expression. (b-e) Single cells having mutant/wild-type Kras (b,c) or Tp53 (d,e) sequencing reads were highlighted in the UMAP.*

**Evolution of TCRs binding to tumor neoantigens from the 344SQ/129Sv mouse model**

Exome-sequencing was performed for the 344SQ/129Sv mice and mutation calling was performed. Mutant peptides generated from tumor somatic mutations were then paired with all mouse MHCs to predict pMHCs *via* netMHCpan/netMHCIIpan. Only predicted strong pMHC binders were retained as putative tumor neoantigens. pMTnet-omni was then exploited to output TCR-pMHC binding predictions by enumerating all possible combinations of pMHCs and TCRs. We subjected these TCR-pMHC pairs to the same analyses as in **Fig. 2d** and **Fig. 2f**. As is shown in **Sup. File 3 Fig. 2** below, conclusions very similar to those from **Fig. 2df** can be drawn.

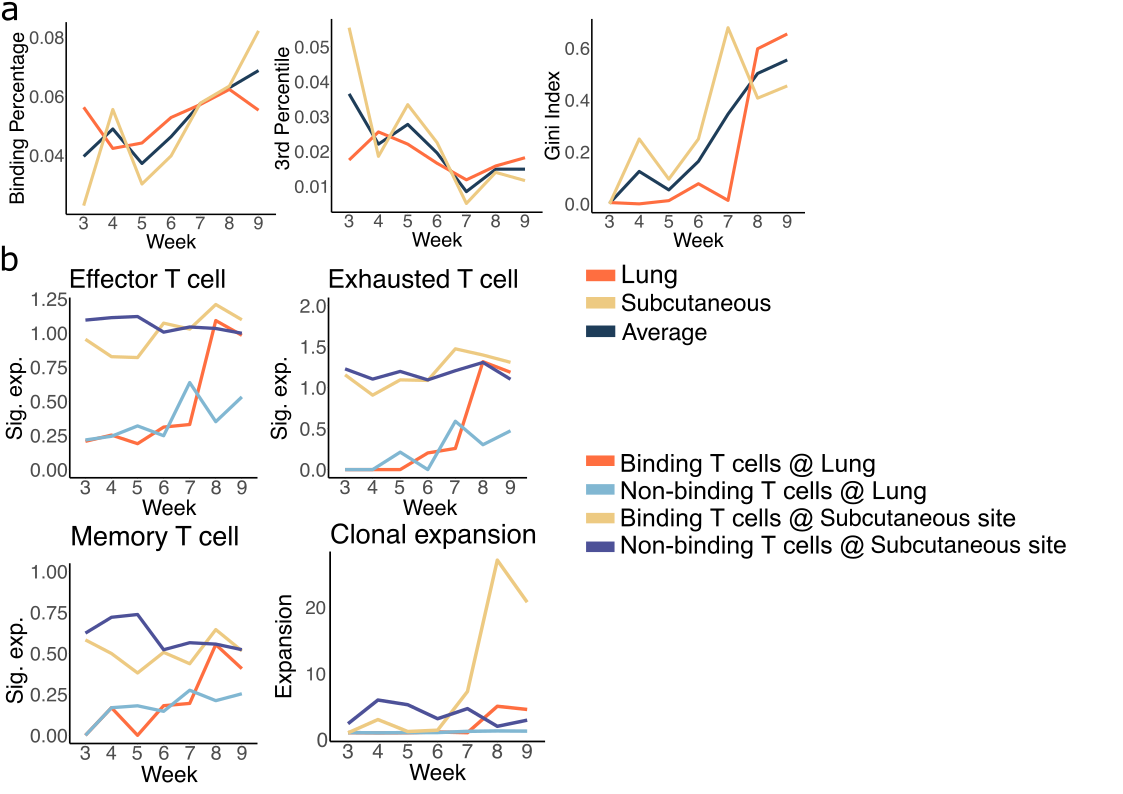

***Sup. File 3 Fig. 2*** *Evolution of TCRs binding to tumor neoantigens from the 344SQ/129Sv mouse model (a) Percentages of predicted binding TCR-pMHC pairs, 3% quantiles of pMTnet-omni TCR-pMHC binding predictions, and Gini indices of the average clonal sizes of TCRs targeting each pMHC, for each week in the subcutaneous tissue samples, lung tissue samples, and their averages. (b) Activation, exhaustion, and memory T cell marker expression and clonal expansions of T cells from the subcutaneous and lung tissue samples for each week.*

**Delineating the spatial heterogeneity of TCR-antigen interactions in a Slide-TCR-seq datasets**

We also analyzed a Slide-TCR-seq dataset from Liu *et al*[*^4^*](https://sciwheel.com/work/citation?ids=13754152&pre=&suf=&sa=0), which was derived from the post-treatment therapy-refractory lung metastasis of a kidney cancer patient. This dataset contains a total of 39,100 sequencing spots with expression data on 25,907 genes and 2,194 unique TCR clonotypes. Cell annotation was performed following marker genes defined by the original study (**Sup. File 3 Fig. 3**). We then used CXCL9, used by Liu *et al*[*^4^*](https://sciwheel.com/work/citation?ids=13754152&pre=&suf=&sa=0) , to define the boundary between tumor epithelial cells and stroma/immune cells. All our following analyses were limited to these sequencing spots near the interaction boundary.

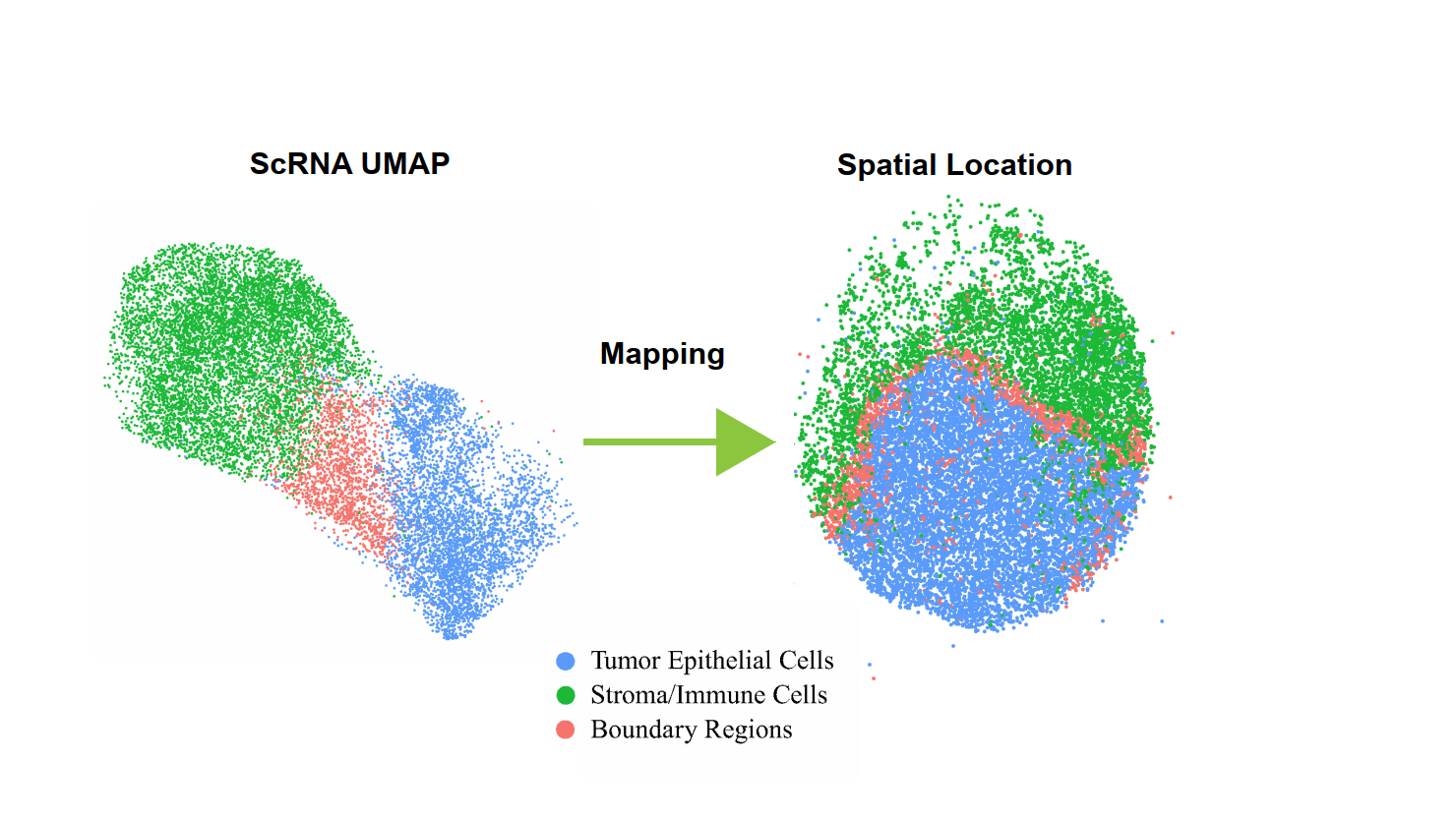

***Sup. File 3 Fig. 3*** *The cell types defined for Slide-TCR-seq sequencing spots in their gene expression and spatial location space.*

Similar to our analyses in **Fig. 5**, we investigated interactions between tumor-associated antigens (TAAs) and T cell receptors. We first defined TAAs by identifying genes that are more highly expressed in the tumor cells as opposed to the stroma/immune cells, and limited our search to genes previously described as putative TAAs (**Sup. Table 3**). Our analyses only resulted in one eligible TAA, *MAGED1*, for this dataset.

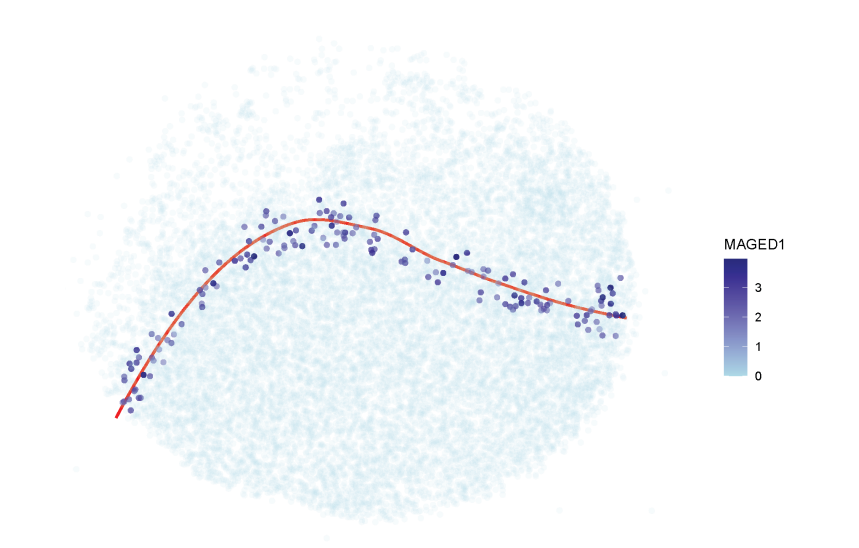

***Sup. File 3 Fig. 4*** *The expression of MAGED1 in the sequencing spots near to the tumor-stroma/immune interaction boundary.*

We showed *MAGED1*’s expression near the interaction boundary (**Sup. File 3 Fig. 4**). As described in **Fig. 5**, we predicted the peptides from this TAA that were presented by this patient’s HLA alleles. Then we use *pMTnet-omni* to predict the interactions between the TAA pMHCs and all TCR clonotypes found in this Slide-TCR-seq dataset. We examined each sequencing spot for the TAA expression, and included the nearest 10 neighbors as well for counting the average clonal expansion size of the TAA-specific TCRs.

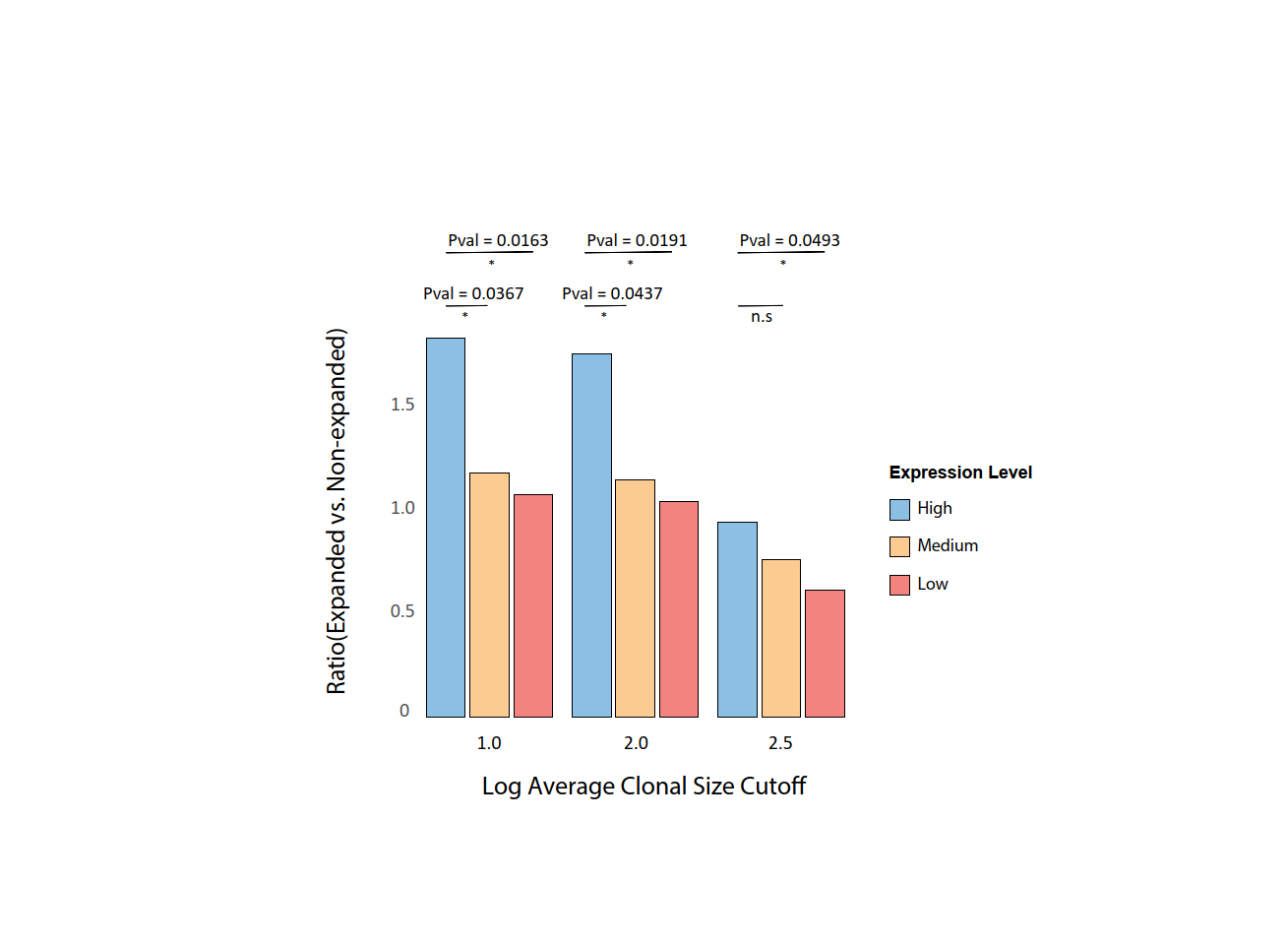

***Sup. File 3 Fig. 5*** *The ratio of the number of sequencing spots with expanded binding TCRs over the number of sequencing with non-expanding binding TCRs, with the logarithm cutoff value to define expanded TCRs shown on the X axis. All sequencing spots near to the interaction boundary was segregated into three subsets based on MAGED1 expression. Chi-squared test was performed and (*) means Pval < 0.05.*

We segregated the sequencing spots into those with high, median and low *MAGED1* expression. As is shown in **Sup. File 3 Fig. 5**, the sequencing spots with high *MAGED1* expression are more likely to be correlated with expanded binding TCR clonotypes**.** Next, we combined the medium and low expressing spots and calculated an Odds Ratio (OR) for *MAGED1*, to investigate whether the high *MAGED1* expressing spots are enriched with expanded binding TCRs, compared with the low/median expressing spots. As in **Fig. 5**, we created a background distribution by random mismatching the TAA-specific TCRs with 5,000 other genes captured in the Slide-TCR-seq dataset. Our results confirmed that the OR of MAGED1 is greater than the average ORs of the background distribution, as shown in **Sup. File 3 Fig. 6.**

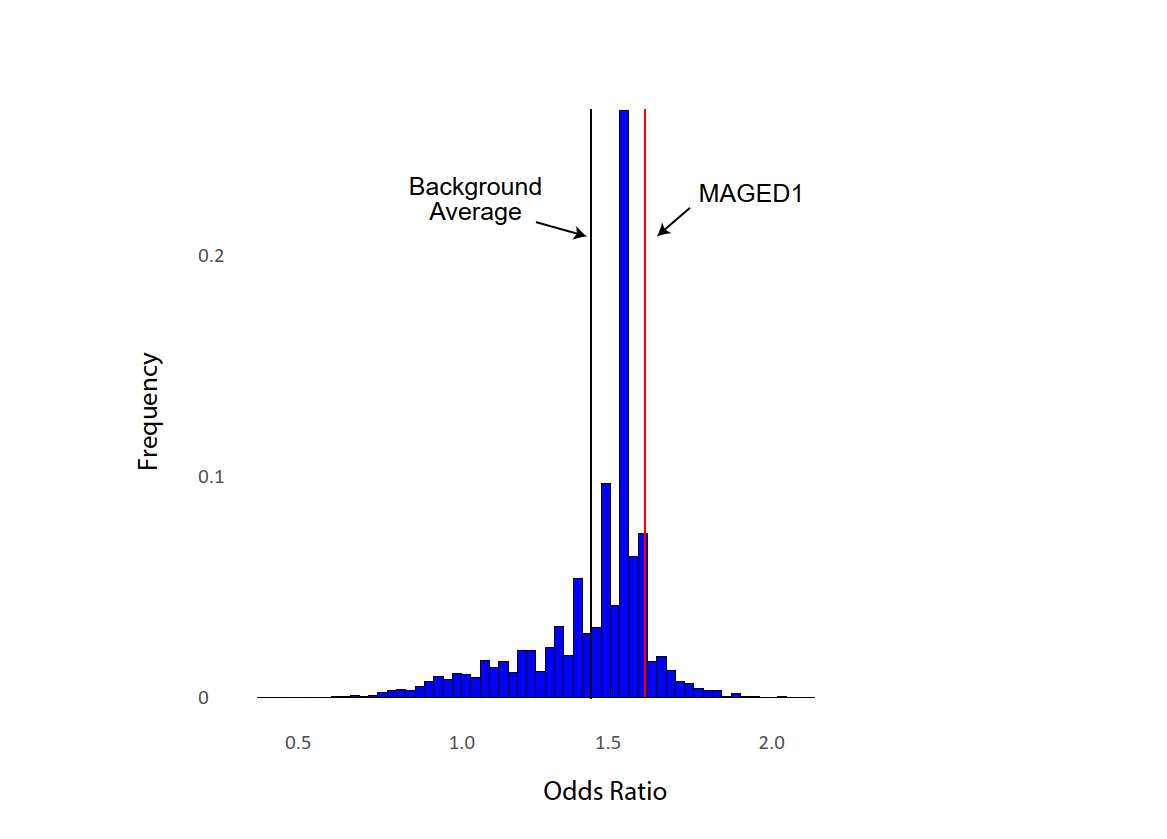

***Sup. File 3 Fig. 6*** *Odds Ratio to test whether the high MAGED1 expressing spots are enriched with expanded binding TCRs, compared with the low/median expressing spots. The red line indicates the OR for MAGED1 and MAGED1 pMHC-specific TCRs, and the black line indicates the average OR of the background distribution by switching MAGED1 with random genes.*

[Bibliography](https://sciwheel.com/work/bibliography)

[1.	Hurskainen, M., Mižíková, I., Cook, D.P., Andersson, N., Cyr-Depauw, C., Lesage, F., Helle, E., Renesme, L., Jankov, R.P., Heikinheimo, M., et al. (2021). Single cell transcriptomic analysis of murine lung development on hyperoxia-induced damage. Nat. Commun. *12*, 1565. 10.1038/s41467-021-21865-2.](https://sciwheel.com/work/bibliography/11169341)

[2.	Maynard, A., McCoach, C.E., Rotow, J.K., Harris, L., Haderk, F., Kerr, D.L., Yu, E.A., Schenk, E.L., Tan, W., Zee, A., et al. (2020). Therapy-Induced Evolution of Human Lung Cancer Revealed by Single-Cell RNA Sequencing. Cell *182*, 1232-1251.e22. 10.1016/j.cell.2020.07.017.](https://sciwheel.com/work/bibliography/9507947)

[3.	Travaglini, K.J., Nabhan, A.N., Penland, L., Sinha, R., Gillich, A., Sit, R.V., Chang, S., Conley, S.D., Mori, Y., Seita, J., et al. (2020). A molecular cell atlas of the human lung from single-cell RNA sequencing. Nature *587*, 619–625. 10.1038/s41586-020-2922-4.](https://sciwheel.com/work/bibliography/10054525)

[4.	Liu, S., Iorgulescu, J.B., Li, S., Borji, M., Barrera-Lopez, I.A., Shanmugam, V., Lyu, H., Morriss, J.W., Garcia, Z.N., Murray, E., et al. (2022). Spatial maps of T cell receptors and transcriptomes reveal distinct immune niches and interactions in the adaptive immune response. Immunity *55*, 1940-1952.e5. 10.1016/j.immuni.2022.09.002.](https://sciwheel.com/work/bibliography/13754152)
